# A trustworthy semantic knowledge network linking data-driven cell phenotypes to diseases and drugs

**DOI:** 10.64898/2026.10.30.763199

**Authors:** Bingfang Xu, Anne Deslattes Mays, Matthew Diller, Raymond LeClair, Ajith V. Pankajam, William Spear, David Osumi-Sutherland, Kumar Arvind, Sreejato Chatterjee, Alexander D. Diehl, Md Mezbahul Islam, Conor Kelly, Rana Khalil, Juhi Khare, Ricardo López Candelaria, Beverly Peng, Latoya McDonald, Daniel Pintard, Noam Rotenberg, David Zweibaum, Yun Zhang, Richard H. Scheuermann

**Author notes:** To whom correspondence should be addressed.,* Name:Richard H. Scheuermann, Address: Division of Intramural Research, National Library of Medicine, National Institutes of Health, Bethesda, MD, USA, Name: Yun Zhang, Address: Division of Intramural Research, National Library of Medicine, National Institutes of Health, Bethesda, MD, USA. co-senior and corresponding authors (Yun Zhang, Richard H. Scheuermann).

## Abstract

Genome-wide association and other genetic studies have identified genes involved in disease pathogenesis, and their encoded proteins have provided therapeutic targets. However, little is known about which cells are the functional home of these disease-associated genes and proteins. Single-cell technologies are enabling high-resolution profiling of cell phenotypes and revealing cellular contexts of disease, yet much of the resulting knowledge remains embedded in individual datasets and scientific literature. To address these gaps, the National Library of Medicine has developed Cell Knowledge Network (NLM-CKN, https://nlm-ckn.org). Reproducible computational workflows have generated cell phenotype knowledge for 2,585 cell sets and 5,139 selectively expressed genes from 88 datasets covering 10 human organs. Mapping of these cell sets to cell types grounds Cell Ontology terms in experimental evidence. This experimental knowledge is integrated with external knowledge of anatomical and cell-type hierarchies, gene-disease associations, and therapeutic interventions. A semantic schema defines node types and relations, enabling knowledge to be transformed into triple assertions and loaded into an ArangoDB database with provenance and quantitative evidence. The NLM-CKN interface supports graphical exploration of the cellular constituents of healthy human tissues. Finally, topological analyses generate insights into cellular contributions to disease pathogenesis, cellular contexts for therapeutic intervention, and drug-repurposing hypotheses.

## Introduction

Cells are the fundamental functional units of life, with different cell types fulfilling different physiological roles in the body. A cell’s identity and function are dictated by the subset of genes and proteins it expresses. The expressed genome defines a distinct cell type, with transitions between cell types occurring during programmed differentiation, while dynamic changes in gene expression in response to environmental cues give rise to transitions between cell states. Together, cell types and cell states determine overall cell phenotypes. Any abnormalities in the expressed genome can contribute to the physical, cellular, and molecular basis of disease. Consequently, understanding normal and abnormal cell phenotypes is key for diagnosing disease and identifying therapeutic targets.

Genome-wide association studies (GWAS) and other genetic approaches have revealed many of the genetic determinants of disease (1–4). However, our understanding of the cellular contributions to disease pathogenesis remains limited. Bridging this gap has become increasingly tractable with the rapid expansion of single-cell technologies, which enable high-resolution profiling of cell phenotypes at unprecedented scale and granularity.

The path to single-cell resolution in the study of cell phenotypes has been gradual. Earlier transcriptomic studies relied on bulk RNA sequencing, which measures average gene expression across cell populations and can therefore mask cellular heterogeneity within tissues. Early single-cell RNA sequencing (scRNA-seq) experiments used low-throughput, predominantly plate-based methods to profile individual cells, providing cellular resolution but with limited scalability (5–8). Single-nucleus RNA sequencing (snRNA-seq) enabled transcriptional profiling of cells in solid organs/tissues in which intact cells are difficult to disperse (9). The advent of droplet-based technologies dramatically increased throughput and has since become the primary engine of single-cell data generation (10–12). In recent years, single-cell multiomics approaches have expanded beyond transcriptomics to profile chromatin accessibility, regulatory states, and protein features (13–15). Meanwhile, spatial transcriptomics reveals information about cellular interactions by measuring gene expression at cellular or near-cellular resolution (16–18).

The rapid accumulation of single-cell data generated by these technologies has necessitated the development of dedicated infrastructure for data reporting, management, and reuse (19,20). Among them, atlas programs and initiatives, such as HuBMAP and the Human Reference Atlas (HRA), provide frameworks for data and metadata standardization (21,22). Data platforms and repositories, including Chan Zuckerberg (CZ) CELLxGENE (23), the Human Cell Atlas (HCA) Data Portal (20), and the Single Cell Expression Atlas (24), as well as more general-purpose repositories such as GEO (25) and ArrayExpress (26), serve as resources for data deposition, sharing, and community analysis. In addition, tissue-specific atlas programs provide organ-specific reference resources. The BICCN/BICAN programs have generated multimodal brain cell atlases (27) with associated data available through NeMO (28), while the Human Lung Cell Atlas (HLCA) (29) and LungMAP CellRef (30) provide reference cell-type classifications for the human respiratory system. The Kidney Precision Medicine Project (KPMP) built a spatially specified human kidney tissue atlas in health and disease with single-cell resolution (31).

Beyond infrastructure for organizing and sharing single-cell data, knowledge derived from the interpretation of these data in biological context is primarily reported as free text in scientific publications. While large language models (LLMs) will play an increasingly important role in extracting knowledge from literature, they are largely limited to findings that authors explicitly describe in the text. Given the massive amount of data generated by a single-cell genomics experiment, only a small fraction of the knowledge that can be derived from a given dataset is summarized in the corresponding publication (32,33). Thus, knowledge bases that transform single-cell data into structured, computable knowledge about cell phenotypes are needed.

Over the past decade, substantial efforts have advanced the development of biomedical knowledge bases in the form of knowledge graphs (KGs), most of which are designed for specific purposes and therefore exhibit different strengths and limitations. Several large-scale KGs that integrate biomedical data from established data sources have been developed to serve specific applications. SPOKE, for example, integrates data from 41 databases and supports precision medicine by connecting patient-specific information with broader networks of biomedical knowledge (34). Developed through the NIH NCATS Biomedical Data Translator program, RTX-KG2 integrates more than 70 biomedical knowledge sources, including UMLS, SemMedDB, ChEMBL, DrugBank, Reactome, and SMPDB, into a KG that supports interoperable access through Translator reasoning systems (35). Complementing such integrated knowledge resources, ROBOKOP supports automated reasoning over distributed biomedical knowledge to address translational questions (36). The Monarch Initiative focuses on genotype-phenotype relationships across species, enabling cross-species reasoning that connects model organism data with human disease (37). PrimeKG (38) is a multimodal biomedical KG that contains clinical features for drugs and diseases and is explicitly designed for precision medicine. The Unified Biomedical Knowledge Graph (UBKG) integrates heterogeneous biomedical knowledge through an ontology-and UMLS-based framework, while Petagraph, built upon UBKG, extends this KG with large-scale genomic and multiomic data (39). Hetionet integrated 29 public resources into a graph containing genes, diseases, compounds, anatomy, pathways, biological processes, molecular functions, side effects, and other entities to predict drug-disease relationships for drug repurposing (40). More recently, approaches for constructing KGs from biomedical literature have also been explored (41,42).

At the infrastructure level, the biomedical community has developed tools for building KGs. For example, the Biolink Model provides a standardized framework for representing biomedical entities and relations and supports ontology-based design of interoperable KGs (43). RTX-KG2 exemplifies the application of the Biolink Model (35). Tools such as PheKnowLator facilitate automated construction of biomedical KGs (44) with an emphasis on FAIR (Findable, Accessible, Interoperable, and Reusable) principles (45), while KG-Hub provides infrastructure for standardized construction, exchange, and reuse of KGs (46).

Despite the advances in biomedical KG development, existing KGs have limited representation of cell phenotypes, especially knowledge derived from single-cell technologies. As a trusted source of data and information from NIH-supported biomedical research programs, the National Library of Medicine (NLM) is uniquely positioned to play a central role in distilling knowledge from research programs and making it broadly accessible. The NLM Cell Knowledge Network (NLM-CKN) was developed to meet this need for systematically capturing and representing knowledge about cell phenotypes derived from single-cell data.

NLM-CKN is a cell-focused biomedical knowledge network with three defining features. First, its core knowledge is derived directly from single-cell experimental data through validated computational pipelines (47–49) that systematically translate sc/snRNA-seq data into knowledge about transcriptional phenotypes in healthy human tissues. Second, this experimentally derived knowledge is integrated with established knowledge about genetic predisposition to disease and therapeutic targets from selected external knowledge resources (50), enabling exploration of the cellular contributions to disease pathogenesis and their potential relevance to drug targeting. Third, the knowledge embedded in the structure and content of OBO Foundry ontologies (51) provides a foundational framework for NLM-CKN.

NLM-CKN has several key characteristics. (1) It is graph-based, representing knowledge as standardized semantically structured (SSS) assertions modeled as subject-predicate-object triples in a graph database. (2) It is interoperable and synergistic, using OBO Foundry ontologies and the Biolink Model (43,51) to facilitate integration with knowledge resources focused on adjacent biomedical domains, while also providing links to external data sources for additional information. (3) It is data-driven, translating experimental data into knowledge at scale through validated computational methods implemented as reproducible modular workflows. (4) It is designed to be trustworthy, with transparent quality control for data selection and the option for user-driven quality filtering. It also provides consistent tracking of knowledge provenance, captures quantitative measures of evidence, and records evidence of reproducibility when the same knowledge is derived from multiple data sources. (5) It is open, exclusively using data sources without restrictive licensing. (6) It is use-case-driven, with nodes and relationships selected to support biological questions—such as identifying cells that express genes implicated in disease—as well as recognizing recurring topological patterns connecting cells, genes, diseases, and therapeutics. (7) It is evolvable, designed for easy expansion with new datasets, knowledge sources, and use cases. (8) It is unique in its cell-centric design, distinguishing it from broader biomedical KGs by organizing knowledge around experimentally derived cell sets and their characterization.

## Materials and Methods

### NLM-CKN infrastructure

The NLM-CKN infrastructure is composed of six major components (Figure 1A). (1) A semantic schema (https://github.com/NIH-NLM/nlm-ckn-schema) based on biomedical ontologies, aligned with the Biolink Model (43), and implemented using the Linked Data Modeling Language (LinkML) (52) serves as the foundation. (2) Single-cell datasets are selected from the CZ CELLxGENE Discover platform (23) based on defined attributes using a custom `cellxgene-harvester-nf` workflow (https://github.com/NIH-NLM/cellxgene-harvester-nf). (3) Cell characteristics are produced using validated computational pipelines, including NS-Forest for marker gene selection (48), implemented through the custom `sc-nsforest-qc-nf` workflow (https://github.com/NIH-NLM/sc-nsforest-qc-nf), which also generates silhouette scores for cell set quality. This is followed by manual mapping of cell sets to Cell Ontology (CL) cell types for reference datasets. NLM-CKN maintains a distinction between cell sets and cell types. Cell types represent standardized classes defined by the CL. Cell sets represent experimentally observed cell populations, often defined by unsupervised clustering of scRNA-seq datasets, that provide evidence for corresponding cell types. (4) External knowledge is retrieved from the Open Targets Platform (50), OBO Foundry ontologies (51), NCBI Gene, UniProt, and other biomedical resources to support disease-related use cases. (5) Subject-predicate-object triple assertions consistent with the NLM-CKN semantic schema, are loaded into an ArangoDB graph database using an extract-transform-load (ETL) process implemented in `nlm-ckn-etl` (https://github.com/NIH-NLM/nlm-ckn-etl). (6) The user interface is designed for graph exploration and workflow development to address specific biomedical use cases (https://github.com/NIH-NLM/nlm-ckn-ui).

**Figure 1.**
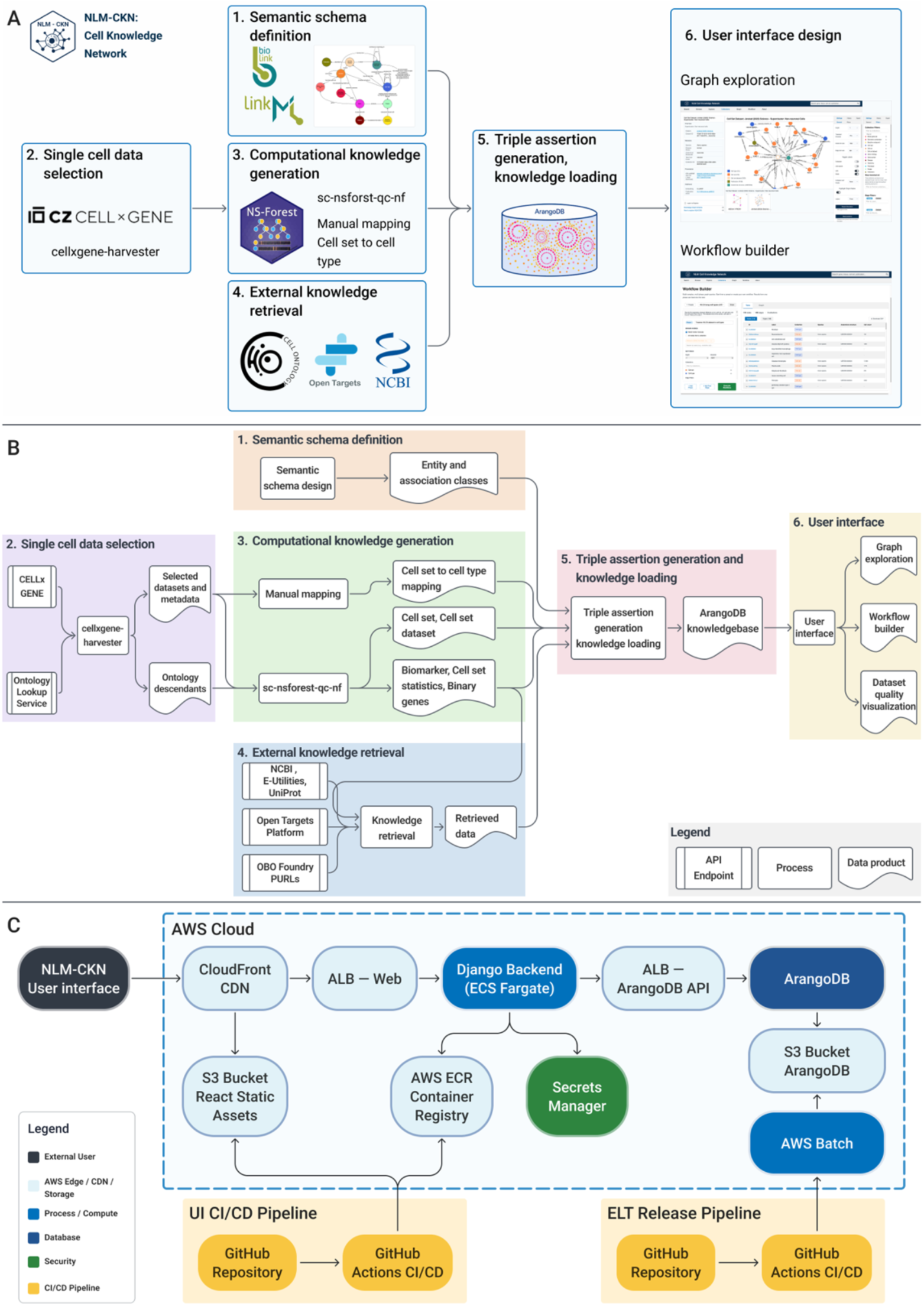
Constructing the NLM Cell Knowledge Network. (**A**) Six major components of the NLM-CKN infrastructure: (1) The NLM-CKN Semantic Schema serves as the underlying representation framework for the knowledge graph and is based on biomedical ontologies, aligned with the Biolink Model, and implemented using LinkML. (2) Single-cell datasets are selected from the CZ CELLxGENE platform using the NLM-CKN cellxgene-harvester-nf workflow. (3) The standardized sc-nsforest-qc-nf computational workflow translates single-cell datasets into structured knowledge about cell set characteristics using the validated NS-Forest computational pipeline. Manual mapping is used to link experimental cell sets to Cell Ontology (CL) cell types for reference datasets. (4) External knowledge is retrieved from the Open Targets Platform, OBO Foundry ontologies, NCBI E-Utilities, and other biomedical resources to support disease-related use cases. (5) Standardized semantically structured subject-predicate-object triple assertions, consistent with the NLM-CKN schema, are generated and loaded into an ArangoDB graph database, integrating knowledge derived from single-cell experiments with external sources of knowledge. (6) A custom graphical user interface allows for faceted and keyword searches, exploratory browsing, and workflow graph query development to address specific biomedical use cases. (**B**) Data processing and information flow of the NLM-CKN: (1) Entity and association classes are defined based on the NLM-CKN Semantic Schema. (2) The cellxgene-harvester-nf workflow accesses the CELLxGENE and Ontology Lookup Service application programming interfaces (APIs) to select single-cell datasets, extract associated metadata, and identify ontology terms and their descendants for experimental variables. (3) Selected datasets are processed using the sc-nsforest-qc-nf workflow, which generates knowledge about cell sets, cell set datasets, biomarkers, binary genes, and cell set statistics. Cell sets from the reference datasets are manually mapped to Cell Ontology (CL) cell types. (4) For genes identified through single-cell data analysis, relevant biomedical knowledge on gene-disease associations, protein-therapeutic compound interactions, and therapeutic compound-disease indications is retrieved from external resources, including NCBI E-Utilities/UniProt, the Open Targets Platform, and OBO Foundry PURLs. (5) Single-cell data products and external knowledge are transformed into schema-conformant SSS triple assertions and loaded into an ArangoDB knowledge graph. (6) The integrated knowledge graph supports a user interface for graph exploration, workflow construction, and dataset quality visualization. (**C**) NLM-CKN deployment architecture. The React-based user interface is distributed through Amazon CloudFront using static assets stored in Amazon S3. Requests to the Django backend are routed through an application load balancer, with the backend deployed as a containerized service on Amazon ECS Fargate. GitHub Actions–based CI/CD workflows support application deployment and ETL releases, with AWS Batch used for ETL processing.

### Use-case-driven design

NLM-CKN employed a use-case-driven design in which use cases were elicited and iteratively refined through collaboration with biologists, bioinformaticians, and clinicians. Each use case addressed one or more of five major categories of inquiry: (1) exploring and evaluating knowledge from single-cell datasets, including their provenance and quality; (2) characterizing the cellular constituents of healthy human tissues, including marker gene information for defined cell populations; (3) identifying gene-disease associations and established or potential therapeutic interventions; (4) linking causal or genetically associated disease genes to their expressing cell sets to investigate cellular contributions to disease pathogenesis; and (5) linking therapeutic target proteins to their expressing cell sets to identify the cellular contexts in which these therapies may exert therapeutic effects. The specific use cases are described in detail in the Results section.

### Semantic schema construction

NLM-CKN was developed using a semantic schema designed to facilitate the integration of knowledge derived from single-cell data with established biomedical knowledge. The overall schema was aligned with the Biolink Model, which is an upper-level data model that standardizes nodes and edges in biomedical KGs. We mapped node types and their relations to corresponding Biolink categories and predicates where applicable. LinkML, a machine-readable schema language, was used to implement the schema and guide KG construction. The NLM-CKN LinkML representation defines node types and semantic relations and specifies the properties associated with nodes and edges (https://github.com/NIH-NLM/nlm-ckn-schema/blob/main/ckn-schema.yaml). These properties include information about data sources, associated experimental evidence, statistical confidence measures, and standard identifiers. Accordingly, data from computational workflows and external sources were structured according to these LinkML-defined node types and relations before being incorporated into the KG. Together, Biolink and LinkML promote semantic harmonization and interoperability with existing biomedical knowledge standards, thereby enabling future integration with other biomedical KG projects.

The NLM-CKN schema was designed to represent both experiment-derived and established biomedical entities as node types. Computational workflows were employed to translate single-cell experimental data into structured representations of cell phenotype knowledge. The experiment-derived node types included Cell_Set, Cell_Set_Dataset, Publication, Biomarker_Combination, and Binary_Gene_Set. The four node types specific to the representation of single-cell experiment-derived knowledge were defined as follows. Cell_Set refers to a collection of cells that share similar molecular profiles and are grouped together, usually by an unsupervised clustering algorithm. Cell_Set_Dataset refers to an assay dataset derived from cells obtained from one or more tissue samples and provides the dataset-level experimental context for its constituent cell sets. Binary_Gene_Set is a collection of genes that are characterized by a “binary expression pattern” (i.e., highly expressed in the target cell set and little to no expression in other cell sets) (47,48). Biomarker_Combination refers to marker gene combinations optimized for cell set classification using metrics that quantify classification confidence (48).

In this study, genes included in a Biomarker_Combination are referred to as marker genes. A marker gene is defined as a gene that is selectively expressed in cells of a given type and can be reliably used alone or in combination as a canonical characteristic to optimally classify them. Genes included in either a Biomarker_Combination or Binary_Gene_Set are collectively referred to as selectively expressed genes, emphasizing their selective expression in the corresponding cell set.

To connect experiment-derived knowledge with established biomedical knowledge and support use cases exploring disease mechanisms and therapeutic interventions, the schema also incorporates Anatomical_Structure, Gene, Cell_Type, Disease, Bioactive_Compound, and Protein node types. Entities represented by these node types were normalized and grounded in established biomedical ontologies and standard bioinformatics representations, including the Cell Ontology (CL) (53), Mondo Disease Ontology (MONDO) (54), Uberon Anatomy Ontology (UBERON) (55), NCBI Gene (56), UniProt (57), and ChEMBL (58).

Knowledge within NLM-CKN was represented as subject-predicate-object triple assertions. Relationships between node types were coded as biologically meaningful edges and mapped to Relation Ontology (RO) (59) object properties and Biolink predicates. Triple assertions were generated from both external biomedical sources and single-cell computational workflows. External assertions included cell type hierarchical relationships from the Cell Ontology, gene-disease and bioactive compound relationships from the Open Targets Platform, anatomical structure hierarchical relationships from UBERON, and gene-protein relationships from UniProt and NCBI Gene. Experiment-derived assertions were generated through CELLxGENE metadata extraction, the NS-Forest pipeline (48), and the FR-Match pipeline (60).

### Computational workflows for single-cell data selection and knowledge generation

NLM-CKN was developed using reproducible computational workflows built with the Nextflow workflow language (61), containerization technologies, and GitHub Actions for continuous integration, version control, and governance. The following sections describe the data sources and selection criteria, cell- and cell set-level filtering, computational workflows for ontology-guided data selection and single-cell knowledge generation, and reference dataset selection.

#### Single-cell data selection criteria

The CZ CELLxGENE Discover resource (23) was used as the primary source of scRNA-seq and snRNA-seq data because it provides several key advantages. It maintains a large and growing collection of single-cell data, aggregating more than 2,240 single-cell datasets comprising over 170 million unique cells and more than 1,180 cell types as of October 18, 2026. CELLxGENE enforces standardized `.h5ad` file formats and harmonized metadata. It has been broadly adopted as a primary data-sharing platform by both individual research laboratories and large atlas consortia, including the BRAIN Initiative (62), the Human Tumor Atlas Network (HTAN) (63), and the Human Cell Atlas (HCA) (32). In addition, CELLxGENE provides a programmatic interface with tools and APIs for data querying and parsing.

NLM-CKN version 1.0 (v1.0) focused on 10 priority healthy human anatomical systems: the respiratory system (UBERON:0001004), retina (UBERON:0000966), brain neocortex (UBERON:0001950), kidney (UBERON:0002113), pancreas (UBERON:0001264), bone marrow (UBERON:0002371), liver (UBERON:0002107), skin of body (UBERON:0002097), heart plus pericardium (UBERON:0015410), and digestive tract (UBERON:0001555). Single-cell datasets for these anatomical structures and their UBERON descendants were queried from CELLxGENE.

Datasets were selected according to the following criteria: (1) only human-derived datasets were included; (2) datasets had to be associated with a peer-reviewed publication; and (3) only scRNA-seq and snRNA-seq datasets were included.

Cells and cell sets within each selected .h5ad dataset file were filtered according to the following criteria: (1) cells not derived from adult tissues or from donors younger than 15 years were excluded; (2) cells not derived from normal tissues of healthy donors were excluded; and (3) cell sets containing fewer than five cells were removed prior to downstream processing.

In addition, derived versions of two datasets were used instead of the versions available in CELLxGENE. For the respiratory system, the CellRef dataset (30), namely, Guo (2023) Nat Commun - “Human CellCards Multi-Study CellRef 1.0 Atlas”, was obtained from LungMAP (https://www.lungmap.net/dataset/?dataset_id=LMEX0000004396).

For the neocortex, the BRAIN Initiative human neocortex data spanning eight cortical areas was selected (62) and the author-processed data was disseminated in five `.h5ad` files in the same CELLxGENE collection (https://cellxgene.cziscience.com/collections/d17249d2-0e6e-4500-abb8-e6c93fa1ac6f). The five datasets are “Supercluster: IT-projecting excitatory neurons”, “Supercluster: MGE-derived interneurons”, “Supercluster: CGE-derived interneurons”, “Supercluster: Non-neuronal cells”, and “Supercluster: Deep layer (non-IT) excitatory neurons”, and the author-provided cross-area cell set annotations were used, to be consistent with the main taxonomy defined in the original publication (62). A global dataset containing sampled cells from all five datasets was used in our workflows and is available for download (Supplementary Table S1).

#### Selection of reference datasets

For each of the 10 priority anatomical systems represented in NLM-CKN v1.0, a single reference Cell_Set_Dataset was selected based on statistical evidence of quality and coverage, including the silhouette score and NS-Forest F-beta score described below, as well as cell-set granularity, cell count per cell set, and community recommendations. All reference datasets are listed in the Reference Dataset collection. Cell sets within each reference cell set dataset were manually curated and mapped to CL cell type terms by comparing author-provided cell set annotations with cell type ontology definitions, marker gene expression patterns, and hierarchical cell type relationships. Mappings were represented using Simple Knowledge Organization System (SKOS) (64) mapping relationships, specifically `skos:exactMatch` and `skos:relatedMatch`.

#### Computational workflows for ontology-guided single-cell dataset selection and processing

We used two computational workflows, `cellxgene-harvester-nf` (https://github.com/NIH-NLM/cellxgene-harvester-nf) and `sc-nsforest-qc-nf` (https://github.com/NIH-NLM/sc-nsforest-qc-nf), in succession for data selection and subsequent knowledge generation. These Nextflow workflows were built with modularity and reproducibility in mind, while leveraging the contributing software packages, including cellxgene-harvester (https://github.com/NIH-NLM/cellxgene-harvester), NS-Forest (https://github.com/NIH-NLM/NSForest), and scsilhouette (https://github.com/NIH-NLM/scsilhouette). Both workflows utilize a publishing step in which the results are published to GitHub. The final filtered `.h5ad` files are published to a publicly accessible Amazon Web Services (AWS) S3 bucket (https://nlm-ckn.nlm.nih.gov/V1.0). For each processed dataset, the workflow also produces a master dataset summary as a JavaScript Object Notation (JSON) file in full data dictionary form. Once published to a branch of the nlm-ckn repository (https://github.com/NIH-NLM/nlm-ckn), reviewed, and merged, the master dataset summary is available for retrieval by the downstream extract–transform–load (ETL) process used to populate the ArangoDB instance.

As shown in Figure 1B, the `cellxgene-harvester-nf` workflow retrieves collections from the CELLxGENE platform and then selects datasets based on ontology annotations. It begins by resolving four ontology-based criteria used for dataset selection and filtering: anatomical structure (UBERON), disease (PATO/MONDO), age (HSAPDV), and assay (EFO). Using the Ontology Lookup Service API, root terms and morphologically related terms for the specified anatomical structure are retrieved from UBERON. Disease terms are then resolved using PATO/MONDO; for NLM-CKN v1.0, the selection was limited to normal tissues. Age terms are resolved using HSAPDV, with v1.0 limited to donors 15 years of age or older. Assays are limited to non-spatial single-cell methods using EFO. The resolved ontology terms are applied during the primary filtering, and qualifying cells are identified based on matches to the relevant UBERON, PATO, HSAPDV, and EFO ontology IDs. `cellxgene-harvester-nf` generates initial filtered `.h5ad` files and records cell counts before and after this primary filtering. Datasets with zero qualifying cells are removed. This primary filtering step does not produce the final filtered dataset, which undergoes additional filtering in the downstream `sc-nsforest-qc-nf` workflow.

The `cellxgene-harvester-nf` workflow fetches the entire dataset collection from CELLxGENE, containing tissue and disease information as structured objects with both labels and ontology term IDs. The collections are then flattened into a dataset-level metadata table, with one row per dataset. Subsequently, publication metadata, including tissue and disease labels and ontology term IDs are generated from the CELLxGENE API response. For each dataset, an additional API call is made to obtain dataset-specific details, including the dataset title, total cell count, `.h5ad` download URL, and CELLxGENE Explorer URL. CELLxGENE release information is also used to determine the GENCODE version for gene symbol annotations; all gene symbols in NLM-CKN v1.0 were obtained from GENCODE release 48. This provenance information enables subsequent findings to be traced to the CELLxGENE release and associated source information. The current process also includes manual inspection and curation, during which the appropriate cell set annotation level and embedding are selected.

#### Computational workflows for single-cell knowledge generation

The resulting datasets from `cellxgene-harvester-nf`, including the initial filtered `.h5ad` files, are used as input to the `sc-nsforest-qc-nf` workflow, which verifies the primary filtering and performs additional filtering, including the removal of clusters containing fewer than five cells. The final filtered datasets are then used for subsequent quality analysis and single-cell knowledge generation. `sc-nsforest-qc-nf` executes NS-Forest and scsilhouette to identify marker genes that distinguish cell sets, compute quantitative metrics, and generate visualizations to aid researchers in evaluating the results. NS-Forest processing includes the generation of original and filtered dendrograms, calculation of cell set statistics, calculation of median expression levels and binary scores of binary genes, batch run of NS-Forest, merging of NS-Forest results, and generation of matrix, dot, and violin plots. Using the same filtered `.h5ad` file as input, the scsilhouette workflow computes cell set-level silhouette scores and produces summary, distribution, and box-plot visualizations. Outputs from both analyses are integrated into combined visualizations and summary statistics at the cell set dataset and cell set levels, including NS-Forest F-beta (beta = 0.5) scores and silhouette scores. All workflow outputs serve as inputs to downstream ETL processes.

#### Workflow reproducibility

`cellxgene-harvester-nf` and `sc-nsforest-qc-nf` are both built using the workflow language Nextflow. Both employ best practices of the elements of style, where each step in the process is a single process (65). Both workflows are modularized so that all steps for any specific package are contained in a single module. Both workflows end with a publication step. This publication step puts the final results of each workflow on a GitHub branch, where they are reviewed before merging. The outputs of these workflows are ingested by the downstream ETL process.

Nextflow allows the same code base to be configured for different computing platforms. Different configurations support continuous integration and testing through GitHub Actions and provide flexibility for users to run the workflows on laptops or in the cloud. Docker is used to build the container images for the production runs. Production runs for this release of NLM-CKN were performed using Lifebit CloudOS, a platform-as-a-service environment, through the NIH STRIDES Initiative on the AWS cloud platform. The flexibility of the workflows also enables derivative projects to run on the NIH on-premises high-performance computing (HPC) system, Biowulf.

All workflow code, configurations, and outputs are managed through the project repository under the NIH-NLM organization on GitHub (https://github.com/NIH-NLM/nlm-ckn), which serves as both a transparency and governance layer for the project. All code is publicly available and fully versioned, with every commit tracked by a unique identifier, timestamp, author, and associated comments, creating a permanent and auditable history of all changes. Workflow outputs are added to the knowledgebase exclusively through a controlled pull request process, ensuring that all changes are reviewed prior to integration. GitHub Actions provide continuous integration and delivery automation for workflow testing and documentation, keeping workflow documentation current as components evolve.

### ETL process for knowledge integration and graph construction

NLM-CKN uses the `nlm-ckn-etl` extract-transform-load process (https://github.com/NIH-NLM/nlm-ckn-etl) to integrate single-cell-derived experimental knowledge with external biomedical knowledge. Both knowledge streams are transformed into triple assertions consistent with the NLM-CKN schema and loaded into the ArangoDB graph (Figure 1B).

#### Single-cell-derived experimental knowledge

The single-cell computational workflows generate knowledge of biomarkers and binary genes, together with experimental context represented by cell sets and cell set datasets (Figure 1B, component 3). In addition, cell sets from the reference datasets are manually mapped to the Cell Ontology cell types. These workflows also generate quantitative quality metrics, including binary scores, F-beta scores, and silhouette scores, which are summarized and visualized at the cell set and cell set dataset levels. This knowledge is transformed into NLM-CKN node types, relations, and associated evidence properties.

#### External knowledge sources

External biomedical knowledge is integrated to connect selectively expressed genes identified from single-cell data with broader molecular, disease, and therapeutic knowledge (Figure 1B, component 4). External knowledge sources were selected based on the following requirements: (1) availability under CC-BY or CC0 licensing; (2) high quality; (3) comprehensiveness of coverage; (4) active maintenance; and (5) use-case-driven selection of knowledge assertions.

The scope of external molecular, disease, and therapeutic knowledge is driven by selectively expressed genes represented in NLM-CKN. Gene nodes are limited to genes selectively expressed in NLM-CKN cell sets through biomarker combinations and binary gene sets; protein nodes are limited to proteins encoded by these genes; disease nodes are limited to diseases associated with these genes with an overall association score greater than 0.5 in the Open Targets Platform; and bioactive compound nodes are limited to compounds that interact with the represented proteins and have reached a minimum “approval” stage for at least one indication in a clinical trial.

#### Ontology processing, graph scoping, and subgraph generation

NLM-CKN represents knowledge as a directed, typed, evidence-based multigraph stored in an ArangoDB database. Biomedical ontologies are used to construct a foundational KG, onto which single-cell-derived experimental knowledge and external biomedical knowledge are subsequently integrated. The foundational KG for NLM-CKN v1.0 was derived from the following ontologies: Cell Ontology (CL), Uberon Anatomy Ontology (UBERON), NCBI organismal taxonomy (NCBITaxon), Mondo Disease Ontology (MONDO), and Protein Ontology (PRO).

Ontologies are downloaded from the OBO Foundr<u>y (</u>http://purl.obolibrary.org/obo/) in OWL format during database construction and parsed using the Apache Jena Ontology API to extract ontology classes, annotations/properties, and relationships. Complex OWL equivalent class axioms are flattened into simpler structures more amenable to subject–predicate–object assertions, with blank nodes replaced by direct edges between entities while retaining provenance from the source ontologies.

Because NLM-CKN v1.0 is scoped to healthy human biology, ontology content is filtered according to taxonomic constraints where applicable. Following integration of single-cell-derived and external knowledge with the foundational KG, a relevant subgraph is generated starting from the Cell_Set nodes. In NLM-CKN v1.0, the complete CL hierarchy was retained, whereas other ontologies were restricted to relevant terms and their ancestor paths. Ancestors were traced to the root using SUB_CLASS_OF relationships for MONDO and PART_OF/CONTRIBUTES_TO_MORPHOLOGY_OF relationships for UBERON. NCBI Taxon was restricted to *Homo sapiens.* The PRO hierarchy was not traversed. This process retained the ontology context required to support NLM-CKN while excluding unrelated ontology content.

#### Schema-guided assertion generation and graph loading

The NLM-CKN LinkML schema guides the transformation of source records into SSS assertions as subject-predicate-object triples. Schema-defined node types and relation classes are implemented as Pydantic classes, which provide an intermediate representation for validating and structuring data prior to graph loading. Python modules are used to retrieve source data and construct subject-predicate-object assertions consistent with the NLM-CKN schema.

The resulting assertions are serialized into an intermediate format and loaded into ArangoDB using a separate Java package. The loading process allows properties such as provenance to be stored directly on relationships when applicable. Other schema-defined properties, including identifiers, experimental context, and quantitative evidence, are retained on their corresponding nodes. The ETL pipeline handles data definition and loading operations, while the backend API uses ArangoDB Query Language (AQL) exclusively for data retrieval to support the user interface. The code for the ETL pipeline is available at https://github.com/NIH-NLM/nlm-ckn-etl.

### User interface design

A web-based user interface was developed to support interactive exploration and querying of NLM-CKN (Figure 1C). The interface is implemented using React for the client-side application, D3.js for interactive visualizations, and the Django REST Framework for communication with the ArangoDB backend. The application is deployed using Amazon Web Services (AWS) cloud infrastructure, with containerized backend services and automated deployment and ETL processes supported through GitHub Actions.

The interface provides three primary approaches for accessing NLM-CKN: faceted browsing, text-based search, and a workflow builder. Faceted browsing allows users to explore knowledge through hierarchical relationships and progressively select entities of interest. Text-based search supports exact, fuzzy (edit-distance), and substring n-gram matching to identify specific entities in the KG. The workflow builder supports multi-step queries by allowing users to define a sequence of graph traversals, with results from one step serving as input to subsequent steps. Predefined workflows corresponding to representative NLM-CKN use cases are provided and can be modified by users.

The interface provides display controls for interactive exploration of query results. Users can filter the displayed graph by entity/collection (node) or relationship (edge) type, expand selected nodes to explore connected knowledge, and remove or retain nodes of interest. Selecting a node displays its associated properties, including identifiers, provenance, and, where applicable, experimental evidence and quantitative statistical confidence. Resulting subgraphs can be exported in SVG, PNG, or JSON format for downstream use. The source code for the user interface is available at https://github.com/NIH-NLM/nlm-ckn-ui.

### Provenance and evidence

NLM-CKN retains provenance and evidence to support the interpretation and evaluation of knowledge assertions. Data-source provenance identifies the origin of each assertion and is recorded as a property of the corresponding edge. For example, CELLxGENE metadata extraction provides structured data used to populate triple assertions connecting cell sets to anatomical structures, cell set datasets, and publications; therefore, as seen in Table 1, CELLxGENE is recorded as the data-source provenance for these assertions. Manual mapping is recorded as the derivation process for the relationships from cell types to cell sets and from cell types to cell set datasets. NS-Forest generates data used to populate knowledge assertions connecting selectively expressed genes with cell sets and is recorded as the source provenance for these assertions. External knowledge assertions similarly retain provenance from their corresponding source resources, including the Open Targets Platform, ontologies such as CL and MONDO, and standard biomedical resources such as NCBI Gene and UniProt. Table 1 records the primary source, the source of process input, and the triple derivation process for each triple assertion type, providing additional provenance details.

**Table 1.**
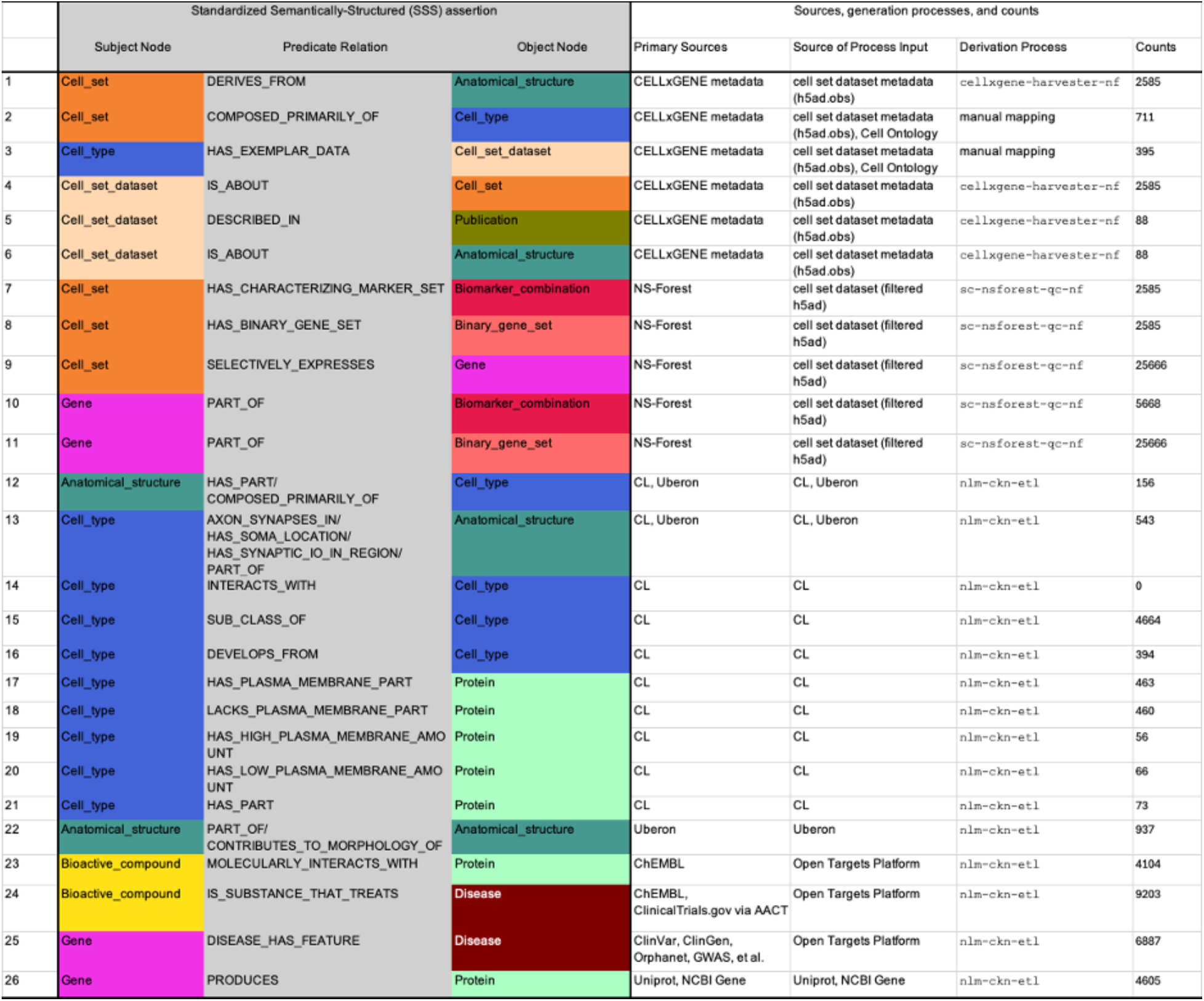
NLM-CKN Standardized Semantically-Structured (SSS) assertion types with their knowledge sources and generation processes.

|  | Standardized Semantically-Structured (SSS) assertion |  |  | Sources, generation processes, and counts |  |  |  |
|---|---|---|---|---|---|---|---|
|  | Subject Node | Predicate Relation | Object Node | Primary Sources | Source of Process Input | Derivation Process | Counts |
| 1 | Cell_set | DERIVES_FROM | Anatomical_structure | CELLxGENE metadata | cell set dataset metadata (h5ad.obs) | cellxgene-harvester-nf | 2585 |
| 2 | Cell_set | COMPOSED_PRIMARILY_OF | Cell_type | CELLxGENE metadata | cell set dataset metadata (h5ad.obs), Cell Ontology | manual mapping | 711 |
| 3 | Cell_type | HAS_EXEMPLAR_DATA | Cell_set_dataset | CELLxGENE metadata | cell set dataset metadata (h5ad.obs), Cell Ontology | manual mapping | 395 |
| 4 | Cell_set_dataset | IS_ABOUT | Cell_set | CELLxGENE metadata | cell set dataset metadata (h5ad.obs) | cellxgene-harvester-nf | 2585 |
| 5 | Cell_set_dataset | DESCRIBED_IN | Publication | CELLxGENE metadata | cell set dataset metadata (h5ad.obs) | cellxgene-harvester-nf | 88 |
| 6 | Cell_set_dataset | IS_ABOUT | Anatomical_structure | CELLxGENE metadata | cell set dataset metadata (h5ad.obs) | cellxgene-harvester-nf | 88 |
| 7 | Cell_set | HAS_CHARACTERIZING_MARKER_SET | Biomarker_combination | NS-Forest | cell set dataset (filtered h5ad) | sc-nsforest-qc-nf | 2585 |
| 8 | Cell_set | HAS_BINARY_GENE_SET | Binary_gene_set | NS-Forest | cell set dataset (filtered h5ad) | sc-nsforest-qc-nf | 2585 |
| 9 | Cell_set | SELECTIVELY_EXPRESSES | Gene | NS-Forest | cell set dataset (filtered h5ad) | sc-nsforest-qc-nf | 25666 |
| 10 | Gene | PART_OF | Biomarker_combination | NS-Forest | cell set dataset (filtered h5ad) | sc-nsforest-qc-nf | 5668 |
| 11 | Gene | PART_OF | Binary_gene_set | NS-Forest | cell set dataset (filtered h5ad) | sc-nsforest-qc-nf | 25666 |
| 12 | Anatomical_structure | HAS_PART/COMPOSED_PRIMARILY_OF | Cell_type | CL, Uberon | CL, Uberon | nlm-ckn-etl | 156 |
| 13 | Cell_type | AXON_SYNAPSES_IN/HAS_SOMA_LOCATION/HAS_SYNAPTIC_IO_IN_REGION/PART_OF | Anatomical_structure | CL, Uberon | CL, Uberon | nlm-ckn-etl | 543 |
| 14 | Cell_type | INTERACTS_WITH | Cell_type | CL | CL | nlm-ckn-etl | 0 |
| 15 | Cell_type | SUB_CLASS_OF | Cell_type | CL | CL | nlm-ckn-etl | 4664 |
| 16 | Cell_type | DEVELOPS_FROM | Cell_type | CL | CL | nlm-ckn-etl | 394 |
| 17 | Cell_type | HAS_PLASMA_MEMBRANE_PART | Protein | CL | CL | nlm-ckn-etl | 463 |
| 18 | Cell_type | LACKS_PLASMA_MEMBRANE_PART | Protein | CL | CL | nlm-ckn-etl | 460 |
| 19 | Cell_type | HAS_HIGH_PLASMA_MEMBRANE_AMOUNT | Protein | CL | CL | nlm-ckn-etl | 56 |
| 20 | Cell_type | HAS_LOW_PLASMA_MEMBRANE_AMOUNT | Protein | CL | CL | nlm-ckn-etl | 66 |
| 21 | Cell_type | HAS_PART | Protein | CL | CL | nlm-ckn-etl | 73 |
| 22 | Anatomical_structure | PART_OF/CONTRIBUTES_TO_MORPHOLOGY_OF | Anatomical_structure | Uberon | Uberon | nlm-ckn-etl | 937 |
| 23 | Bioactive_compound | MOLECULARLY_INTERACTS_WITH | Protein | ChEMBL | Open Targets Platform | nlm-ckn-etl | 4104 |
| 24 | Bioactive_compound | IS_SUBSTANCE_THAT_TREATS | Disease | ChEMBL, ClinicalTrials.gov via AACT | Open Targets Platform | nlm-ckn-etl | 9203 |
| 25 | Gene | DISEASE_HAS_FEATURE | Disease | ClinVar, ClinGen, Orphanet, GWAS, et al. | Open Targets Platform | nlm-ckn-etl | 6887 |
| 26 | Gene | PRODUCES | Protein | Uniprot, NCBI Gene | Uniprot, NCBI Gene | nlm-ckn-etl | 4605 |

NLM-CKN also retains experimental evidence supporting single-cell knowledge. For example, cell sets, defined as groups of cells often identified by unsupervised clustering of single-cell transcriptomic data, provide experimental evidence for the corresponding cell types. For the reference datasets, cell sets were manually mapped to Cell Ontology (CL) terms through the COMPOSED_PRIMARILY_OF relation, allowing each mapped cell type to be traced to its supporting experimental cell set. Cell sets were further linked to the cell set datasets from which they were derived, preserving experimental context and enabling cell type knowledge to be traced back to the underlying experiments.

NLM-CKN also records quantitative statistical confidence measures generated by the computational pipelines as evidence. The NS-Forest F-beta score measures the ability of a biomarker combination to distinguish a target cell set from other cell sets within a cell set dataset and therefore provides quantitative confidence in the biological separability of cell sets. Because the F-beta score reflects biological separability based on marker genes, it captures both cell set clustering quality and biomarker combination classification performance and is recorded as a property of the corresponding Cell_Set and Biomarker_Combination nodes. At the dataset level, the median F-beta score across all Biomarker_Combination nodes within a dataset is recorded as a property of the corresponding Cell_Set_Dataset node.

The binary score, also generated by the NS-Forest pipeline, measures the expression specificity of individual genes for a target cell set. The mean binary score across all genes in a Binary_Gene_Set is recorded as a property of the corresponding Binary_Gene_Set node, providing a summary measure of gene expression specificity for that binary gene set.

Finally, the scsilhouette pipeline, a component of `sc-nsforest-qc-nf`, calculates silhouette scores to assess the compactness and separation of cell sets in the author-provided reduced-dimensional space (e.g., UMAP). The median silhouette score for each cell set is recorded as a property of the corresponding Cell_Set node, while the median of the cell set-level median silhouette scores is recorded as a property of the corresponding Cell_Set_Dataset node. The quantitative evidence associated with each node type is summarized in Table 2.

**Table 2.** NLM-CKN node types with their associated quantitative evidence and properties.

| Node | Node type | Node ID sources | Source origin | Counts | Evidence | Node property display |
|---|---|---|---|---|---|---|
| 1 | Biomarker_combination | internally generated identifiers:BMC/XXXXXX | NS-Forest pipeline | 2585 | NS-Forest F-beta score | Markers & Selectively Expressed Genes, Biomarker Combination Metrics |
| 2 | Binary_gene_set | internally generated identifiers: BGS/XXXXXX | NS-Forest pipeline | 2585 | median of binary score | Markers & Selectively Expressed Genes, Binary Score |
| 3 | Cell_set | internally generated identifiers: CS/XXXXXX | cellxgene metadata | 2585 | Silhouette score,NS-Forest F-beta score | Context, Provenance, Analysis Metadata, CKN Filtering Criteria, Post-filtering Cell Set Metadata, Post-filtering Cell Set Statistics, Markers & Selectively Expressed Genes, Biomarker Combination Metrics, Additional |
| 4 | Cell_set_dataset | internally generated identifiers: CSD/XXXXXX | cellxgene metadata | 88 | median F-beta score, median of median Silhouette score | Context, Provenance, Analysis Metadata, CKN Filtering Criteria, Post-filtering Dataset Metadata, Post-filtering Dataset Statistics, Additional |
| 5 | Bioactive_compound | ChEMBL | https://www.ebi.ac.uk/chembl/ | 1957 |  | Label, Protein, UniProt ID, Drug type, Definition, Mechanism of action, Trade names, Exact synonym(s), Approval status |
| 6 | Cell_type | CL | http://purl.obolibrary.org/obo/cl.obo | 3337 |  | Label, Definition, DB cross reference(s) |
| 7 | Publication | DOI | http://doi.org/ | 60 |  | Publication, Authors, Year, Title, Journal |
| 8 | Disease | MONDO | http://purl.obolibrary.org/obo/mondo.obo | 5159 |  | Label, Definition, Exact synonym(s), DB cross reference(s) |
| 9 | Gene | NCBI_Gene | https://www.ncbi.nlm.nih.gov/gene | 5139 |  | Gene symbol, Gene id, Exact synonym(s), Gene type, Label, RefSeq summary, Reference sequence identifier, mRNA (NM) and protein(NP) sequences, Species |
| 10 | Anatomical_structure | UBERON | http://purl.obolibrary.org/obo/uberon.obo | 1028 |  | Label, Definition, Exact synonym(s), DB cross reference(s) |
| 11 | Protein | UniProt | https://www.uniprot.org/ | 4822 |  | Protein name, UniProt ID, Number of amino acids, Protein function, Species, Gene symbol, Annotation score, Label, Definition, OBO namespace, Exact synonym, DB cross reference(s) |

The conceptual framework for ensuring trustworthiness through provenance and evidence in NLM-CKN, including cell sets as experimental evidence for cell types and quantitative assessment of cell set quality, has been described in detail elsewhere (66).

## Results

### NLM-CKN v1.0 schema

Figure 2 provides a graphical representation of the NLM-CKN v1.0 schema, illustrating the node types and their relationships as graph triples. A line separates node types derived from single-cell experiments from external knowledge node types. The four experimental node types—Cell_Set, Cell_Set_Dataset, Binary_Gene_Set, and Biomarker_Combination—are connected to node types grounded in ontologies and standard biological representations, including Cell_Type, Anatomical_Structure, and Gene. Among these relationships, two triples are particularly important in NLM-CKN. The Cell_Set-COMPOSED_PRIMARILY_OF-Cell_Type triple maps experimentally derived cell sets to Cell Ontology cell type terms, grounding experimental observations within a standardized ontological framework. The Cell_Set-SELECTIVELY_EXPRESSES-Gene triple connects single-cell-derived knowledge with established knowledge of genetic contributions to disease and therapeutic interventions, providing cellular context for disease-associated gene expression and enabling exploration of cellular contributions to disease mechanisms and therapeutic targets.

**Figure 2.**
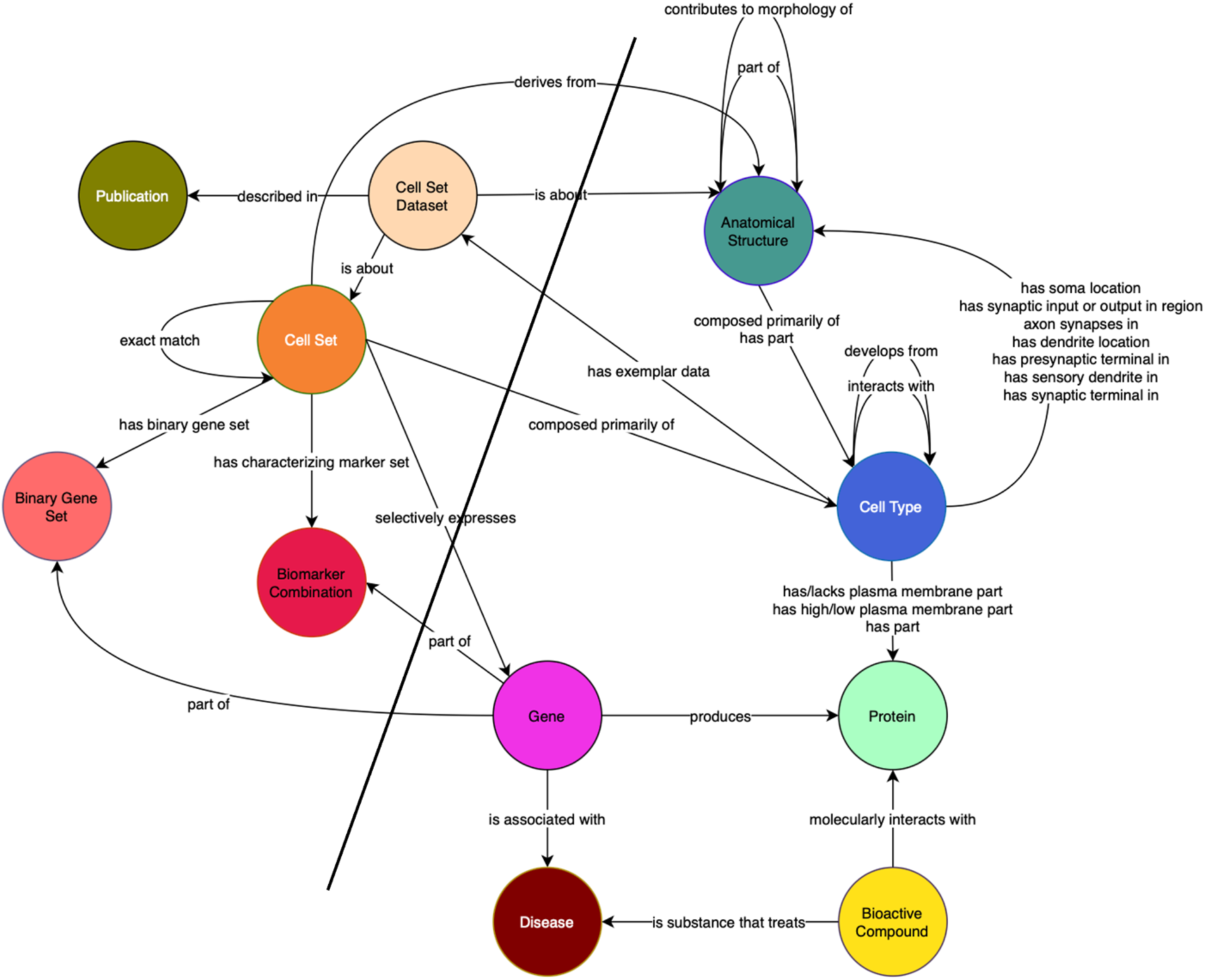
Schematic representation of the NLM-CKN schema. Node types and relations derived from single-cell experiments are shown to the left of the blue line. Four experimental node types (Cell Set Dataset, Cell Set, Binary Gene Set, and Biomarker Combination) are directly connected to external knowledge node types, including Cell Type, Anatomical Structure, and Gene, which are further connected to Disease, Protein, and Bioactive Compound node types. Edges describe the semantic relations between node types. Node types and relations are described in detail in Tables 1 and 2.

Table 2 summarizes 11 node types in NLM-CKN v1.0, including their node ID sources, origin sources, counts, associated evidence metrics, and node properties. Rows 1-4 of Table 2 describe four empirical node types derived from single-cell experiments, where each node was assigned a unique, internally generated identifier. Cell_Set is the central empirical node type. Each Cell_Set is associated with the corresponding Biomarker_Combination and Binary_Gene_Set nodes and links to a Cell_Set_Dataset derived from an Anatomical_Structure described in a Publication. Rows 5-11 of Table 2 show seven node types that use IDs from established ontologies or standard bioinformatics representations, consistent with conventions used in other biomedical databases.

Biomarker_Combination and Binary_Gene_Set nodes were generated by the NS-Forest pipeline. The “Evidence metric” column in Table 2 provides the quantitative confidence measures computed by the computational pipelines. Biomarker_Combination nodes store the NS-Forest F-beta score. Binary_Gene_Set nodes store the mean binary score. Cell_Set and Cell_Set_Dataset nodes were established through CELLxGENE metadata extraction. Silhouette scores and NS-Forest F-beta scores reflect the quality of cell sets, whereas the median F-beta score and median of median silhouette score represent Cell_Set_Dataset-level summary statistics.

The “Node property display” column in Table 2 summarizes the properties displayed for each node type in the NLM-CKN interface, including node context and metadata, evidence metrics, provenance information with links to the original resources, and additional supporting information. These properties are displayed in the left panel of each node landing page in the NLM-CKN user interface.

Table 1 summarizes the 26 standardized semantically-structured (SSS) assertion types in NLM-CKN v1.0, including their subject nodes, predicate relations, object nodes, primary sources, source of process input, derivation process, and counts. Rows 1-11 show 11 SSS assertion types created from single-cell experimental data through computational pipelines. Rows 1-6 show six SSS assertion types originating from CELLxGENE metadata extraction. Rows 7-11 show five SSS assertion types generated by the NS-Forest pipeline. Rows 12-26 show 15 SSS assertion types derived from external data sources representing established biological knowledge. Rows 12-21 show ten SSS assertion types mainly extracted from the Cell Ontology, capturing hierarchical relationships among cell types. Hierarchical relationships among anatomical structures were obtained from UBERON (row 22), while SSS assertions linking genes, diseases, and bioactive compounds were sourced from the Open Targets Platform (rows 23–25). Gene-PRODUCTS-Protein relationships were drawn from UniProt and NCBI Gene (row 26).

### Use cases and workflow builder

Table 3 presents the 12 use cases that guided the development of NLM-CKN. These use cases are implemented as predefined queries in the Workflow Builder on the NLM-CKN website (https://nlm-ckn.org/#/workflow-builder), where the highlighted search terms (bold) can be replaced with user-supplied terms to modify the predefined queries. Use Case #1 queries single-cell datasets and their associated quality evidence. Use Cases #2-5 query single-cell-derived knowledge and Cell Ontology knowledge to characterize cell phenotypes and cellular constituents of healthy human tissues. Use Cases #6-12 extend these queries by connecting cell set/type-selective gene expression with disease and therapeutic knowledge from external biomedical resources, demonstrating how NLM-CKN supports investigation of cellular contributions to disease mechanisms and the cellular context of therapeutic targets.

**Table 3.**
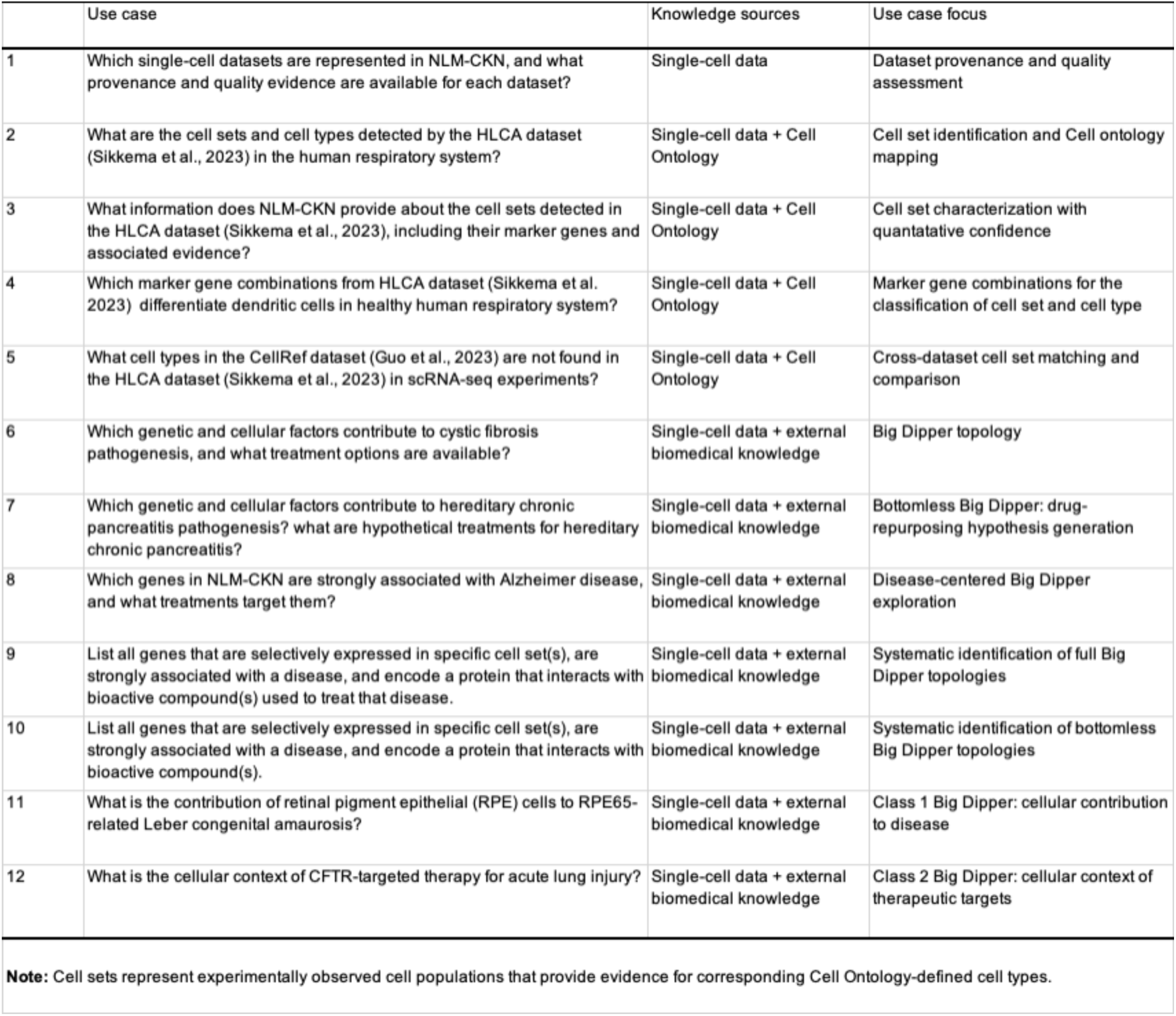
Use case questions, focus, and knowledge sources.

|  | Use case | Knowledge sources | Use case focus |
|---|---|---|---|
| 1 | Which single-cell datasets are represented in NLM-CKN, and what provenance and quality evidence are available for each dataset? | Single-cell data | Dataset provenance and quality assessment |
| 2 | What are the cell sets and cell types detected by the HLCA dataset (Sikkema et al., 2023) in the human respiratory system? | Single-cell data + Cell Ontology | Cell set identification and Cell ontology mapping |
| 3 | What information does NLM-CKN provide about the cell sets detected in the HLCA dataset (Sikkema et al., 2023), including their marker genes and associated evidence? | Single-cell data + Cell Ontology | Cell set characterization with quantatative confidence |
| 4 | Which marker gene combinations from HLCA dataset (Sikkema et al. 2023) differentiate dendritic cells in healthy human respiratory system? | Single-cell data + Cell Ontology | Marker gene combinations for the classification of cell set and cell type |
| 5 | What cell types in the CellRef dataset (Guo et al., 2023) are not found in the HLCA dataset (Sikkema et al., 2023) in scRNA-seq experiments? | Single-cell data + Cell Ontology | Cross-dataset cell set matching and comparison |
| 6 | Which genetic and cellular factors contribute to cystic fibrosis pathogenesis, and what treatment options are available? | Single-cell data + external biomedical knowledge | Big Dipper topology |
| 7 | Which genetic and cellular factors contribute to hereditary chronic pancreatitis pathogenesis? what are hypothetical treatments for hereditary chronic pancreatitis? | Single-cell data + external biomedical knowledge | Bottomless Big Dipper: drug-repurposing hypothesis generation |
| 8 | Which genes in NLM-CKN are strongly associated with Alzheimer disease, and what treatments target them? | Single-cell data + external biomedical knowledge | Disease-centered Big Dipper exploration |
| 9 | List all genes that are selectively expressed in specific cell set(s), are strongly associated with a disease, and encode a protein that interacts with bioactive compound(s) used to treat that disease. | Single-cell data + external biomedical knowledge | Systematic identification of full Big Dipper topologies |
| 10 | List all genes that are selectively expressed in specific cell set(s), are strongly associated with a disease, and encode a protein that interacts with bioactive compound(s). | Single-cell data + external biomedical knowledge | Systematic identification of bottomless Big Dipper topologies |
| 11 | What is the contribution of retinal pigment epithelial (RPE) cells to RPE65-related Leber congenital amaurosis? | Single-cell data + external biomedical knowledge | Class 1 Big Dipper: cellular contribution to disease |
| 12 | What is the cellular context of CFTR-targeted therapy for acute lung injury? | Single-cell data + external biomedical knowledge | Class 2 Big Dipper: cellular context of therapeutic targets |

**Note:** Cell sets represent experimentally observed cell populations that provide evidence for corresponding Cell Ontology-defined cell types.

### Use cases demonstrating the qualities of single-cell datasets

Use Case #1 addresses the question: “Which single-cell datasets are represented in NLM-CKN, and what provenance and quality evidence are available for each dataset?”. NLM-CKN v1.0 includes knowledge from 88 single-cell datasets. Supplementary Table S1 provides a comprehensive summary of key provenance and quality evidence for all Cell_Set_Dataset nodes, including two summary metrics reflecting dataset quality: the median of the median silhouette score and the median NS-Forest F-beta score across biomarker combinations. Supplementary Table S1 also reports cell counts before and after filtering, the total number of remaining cell sets, and metadata, including original publication DOIs, direct links to the CELLxGENE Explorer, and links to download the filtered dataset `.h5ad` files. Together, this information allows users to evaluate dataset quality and trace each dataset to its original source.

To enable users to visualize the quality of each dataset, box-and-whisker plots of silhouette scores together with bar plots of F-beta scores for marker gene combinations across all cell sets within each dataset are stored as properties of the corresponding Cell_Set_Dataset nodes in NLM-CKN. The plots are displayed below the main dataset subgraph. Supplementary Figure S1 shows the combined metrics plot for the pancreas reference dataset (Xu (2023) Cell – “Pancreas” for pancreas) (67).

Dendrograms, violin plots, and dot plots are displayed for each Cell_Set_Dataset to provide users with additional dataset information. For example, Supplementary Figure S2 shows a dendrogram for the pancreas reference dataset, generated by hierarchical clustering based on similarity in the transcriptional profiles, illustrating the hierarchical relationships between cell sets. Supplementary Figure S3-S4 show a violin plot and a dot plot of NS-Forest marker gene expression for each cell set in the same dataset, respectively, illustrating the specificity of the marker gene combinations.

### Use cases demonstrating anchoring of Cell Ontology terms in experimental evidence

Use Case #2 investigates the question: “What are the cell types detected in the HLCA dataset (Sikkema et al., 2023) from the human respiratory system?” The HLCA dataset, with the unique dataset name “Sikkema (2023) Nat Med – ‘An integrated cell atlas of the human lung in health and disease (core)’ for respiratory system”, serves as the reference dataset for the human respiratory system and is used to demonstrate several use cases.

As shown in Figure 3, the human respiratory system, an anatomical structure node, is linked to the HLCA Cell_Set_Dataset node through the IS_ABOUT relation. The HLCA dataset contains 61 cell sets (orange nodes) mapped to 57 cell types (blue nodes) through the COMPOSED_PRIMARILY_OF relation. The expanded subgraph illustrates different SKOS mappings between cell sets and cell types, including 1-to-1 `skos:exactMatch` and many-to-1 `skos:relatedMatch` mappings. For example, three smooth muscle cell sets map to a single cell type using `skos:relatedMatch`. In addition, the graph captures the hierarchical organization of CL cell types through the SUB_CLASS_OF relation, enabling exploration of cell type classification within the human respiratory system. This use case demonstrates how NLM-CKN links cell sets identified by single-cell experiments to the hierarchical structure of the Cell Ontology.

**Figure 3.**
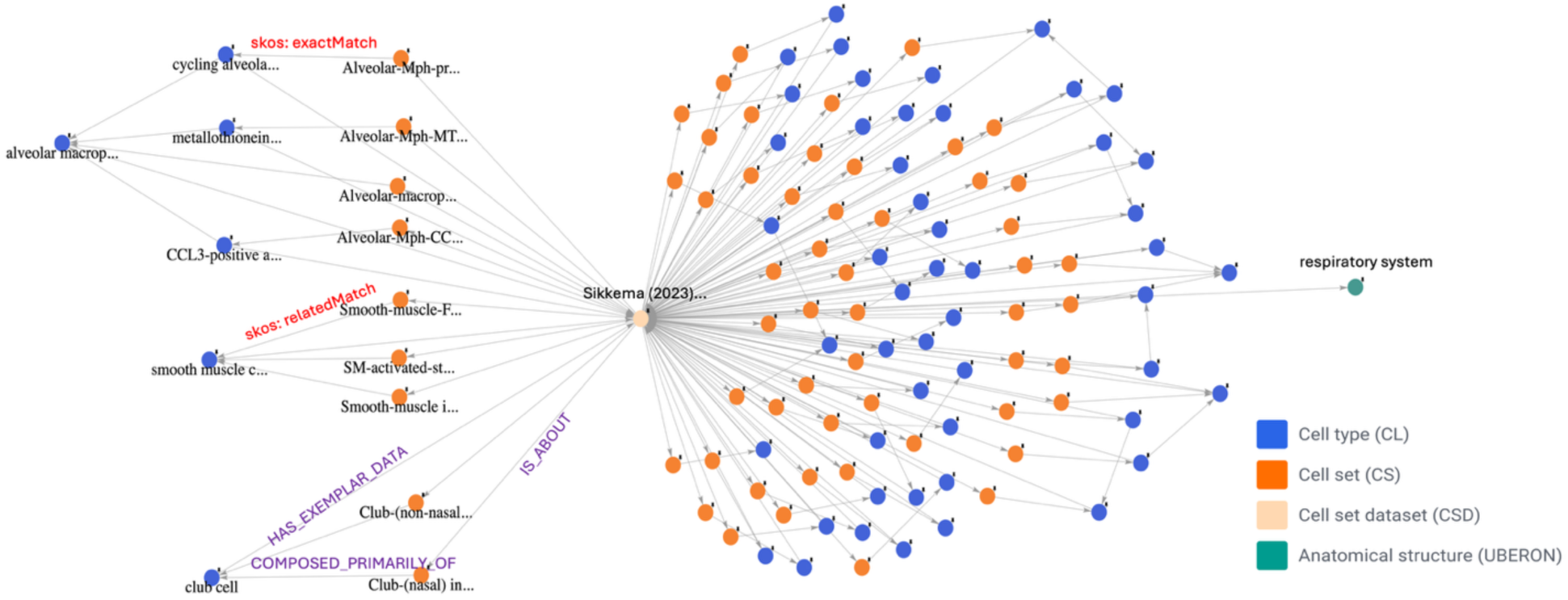
Identification of cell types in the human respiratory system (Use Case #2). The Sikkema 2023 HLCA cell set dataset (tan node) contains 61 cell sets (orange nodes) mapped to 57 Cell Ontology (CL) cell types (blue nodes) in the human respiratory system (green node), with the cell set dataset linked to cell sets with IS_ABOUT relations, cell types linked to the cell set dataset node with HAS_EXEMPLAR_DATA relations, and cell types linked to cell sets with COMPOSED_PRIMARILY_OF relations. The expanded subgraph on the left illustrates different mappings between cell sets and CL cell types, including 1:1 `skos:exactMatch` and 2:1 and 3:1 `skos:relatedMatch` mappings, highlighting gaps in the hierarchical organization of cell types within the Cell Ontology.

In addition to the subgraph view illustrated in Figure 3, the workflow builder provides a tabular view summarizing the properties of each node. This tabular view directly answers Use Case #3: “What are the marker genes, cell count, and other information for the cell types detected in the HLCA dataset (Sikkema et al., 2023) from the human respiratory system?”. Supplementary Table S2 presents the 61 cell set nodes (`author_cell_term`) identified from the HLCA dataset, together with their associated properties, including binary gene set, biomarker combination, cell count, F-beta score of biomarker combination, median silhouette score, ontology PURL, cell type name, and publication.

Use Case #4 asks: “Which marker gene combinations from the HLCA dataset (Sikkema et al., 2023) differentiate dendritic cell types in the healthy human respiratory system?” As shown in Figure 4, dendritic cell, a cell type node, is connected to four dendritic cell subtypes through direct or multi-step SUB_CLASS_OF relationships. These four cell types are associated with four cell sets from the HLCA via the COMPOSED_PRIMARILY_OF relationship. Each cell set is connected to its own biomarker combination through the HAS_CHARACTERIZING_MARKER_SET relationship, with individual genes linked to the biomarker combination via the PART_OF relationship. Therefore, *CCL22, JCHAIN, LILRA4, HLA-DQA1, CLEC10A, CLEC9A*, and *DNASE1L3* are the marker genes that differentiate the four dendritic cell types: lung migratory dendritic cell, conventional dendritic cell, plasmacytoid dendritic cell, and CD1c-positive myeloid dendritic cell in the human respiratory system. Use Case #4 highlights the value of integrating single-cell data with the hierarchical structure of the Cell Ontology.

**Figure 4.**
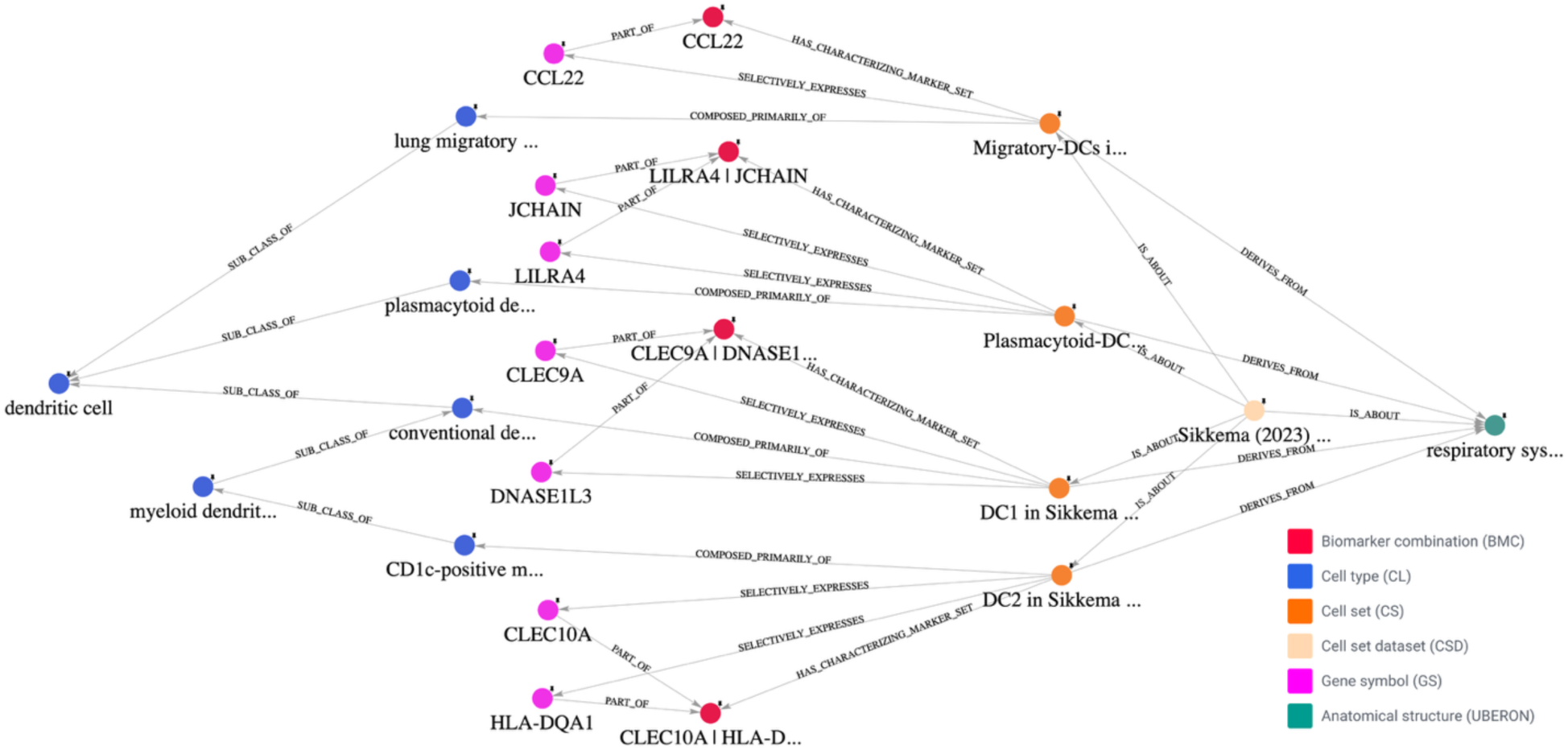
Marker gene combinations that differentiate dendritic cell types in the healthy human respiratory system (Use Case #4). Four cell sets (orange nodes) from the Sikkema 2023 HLCA dataset (tan node) are mapped to dendritic cell subtypes in the Cell Ontology (CL) (blue nodes). Each cell set is characterized by a unique biomarker combination (red nodes) linked to its constituent marker genes (pink nodes). These marker gene combinations distinguish the four dendritic cell sets/types in the human respiratory system (green node).

Use Case #5 addresses the question: “Which cell types represented in the CellRef dataset (Guo et al., 2023) are not represented in the HLCA dataset (Sikkema et al., 2023)?” The HLCA dataset serves as the reference dataset for the respiratory system. A total of 61 cell sets from HLCA were manually mapped to cell types in the Cell Ontology. 42 out of 48 cell sets from CellRef were matched to HLCA cell sets using FR-Match and other algorithms (60). When a match was identified, the CellRef cell set was connected to the cell type node already linked to the corresponding HLCA cell set in the graph.

As shown in Figure 5, the cell set-cell type pairs are circled in different colors to reflect their dataset membership. The pairs in red circles represent pairs shared by CellRef and HLCA, indicating cell sets that match between datasets. The pairs in green circles are unique to HLCA, indicating cell sets/types present in the HLCA dataset but not found in CellRef. The pairs in blue circles represent the six cell sets from CellRef that did not have matches in HLCA and are therefore unique to CellRef. These six cell sets were subsequently manually mapped to six Cell Ontology terms.

**Figure 5.**
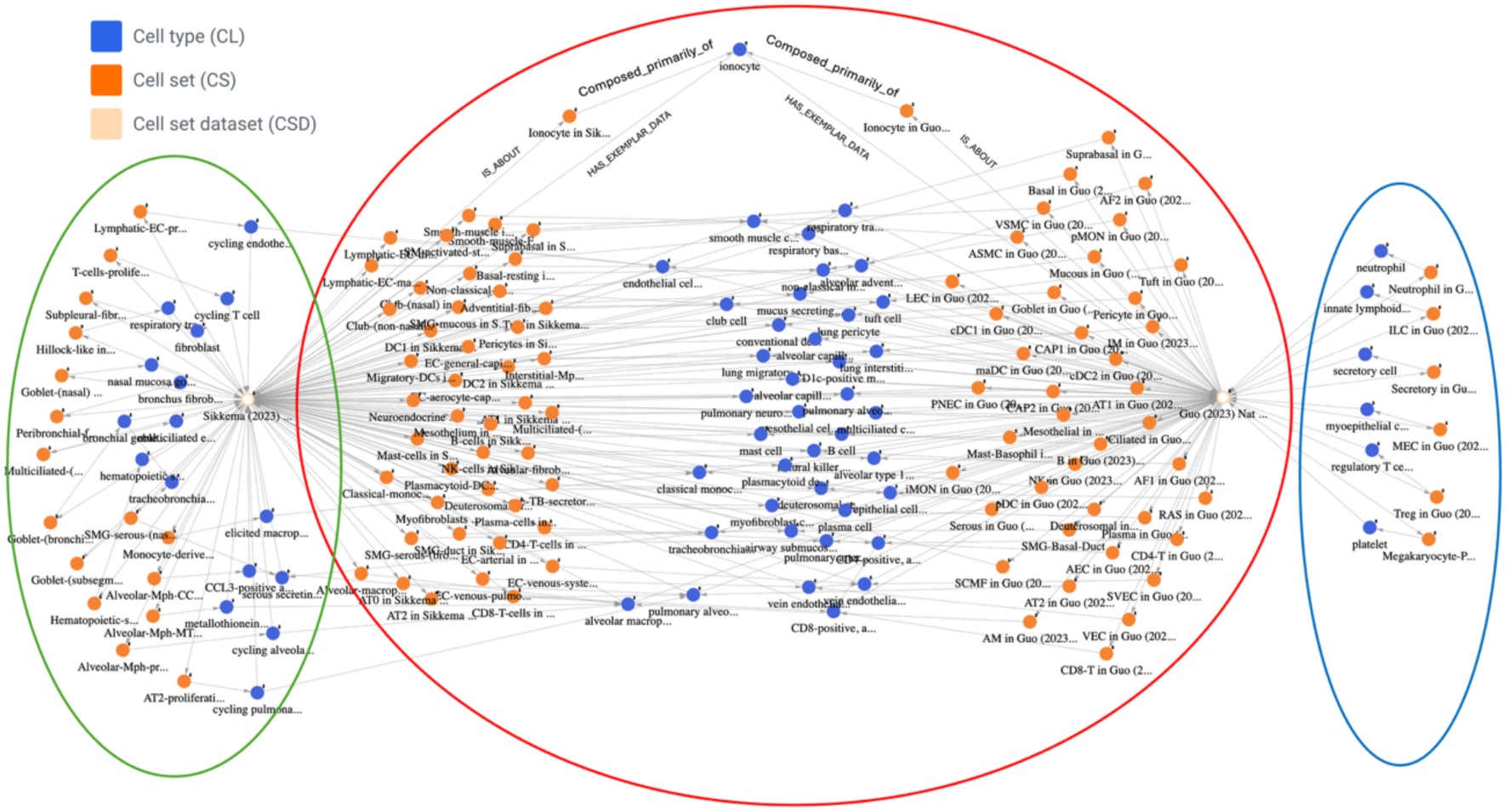
Identification of shared and specific cell sets in the CellRef and HLCA datasets (Use Case #5). FR-Match identified 42 matching cell set pairs shared between the reference Sikkema 2023 HLCA dataset (61 cell sets) and the Guo 2023 CellRef dataset (48 cell sets), enabling the transfer of corresponding Cell Ontology (CL) cell type annotations to matched CellRef cell sets. Matched cell sets are highlighted within the red oval, HLCA-specific cell sets within the green oval, and CellRef-specific cell sets within the blue oval. The expanded subgraph at the top shows that the HLCA and CellRef ionocyte cell sets are both connected to the ionocyte CL cell type with COMPOSED_PRIMARILY_OF relations.

### Big Dipper topology for exploring disease mechanisms and therapeutic interventions

#### Big Dipper

When exploring use cases involving disease mechanisms and therapeutic interventions, we identified a topological structure in NLM-CKN in which seven node types can be arranged in a shape that resembles the Big Dipper stellar constellation, as shown in Figure 6A. In this topology, a selectively expressed gene is associated with a disease, and the protein produced by the gene molecularly interacts with a bioactive compound that treats the disease. Simultaneously, the gene is selectively expressed in cell sets that belong to a cell set dataset derived from a specific anatomical structure. Therefore, the Big Dipper topology captures not only the gene-disease association and therapeutic intervention but also the anatomical and cellular expression patterns of disease-associated genes.

**Figure 6.**
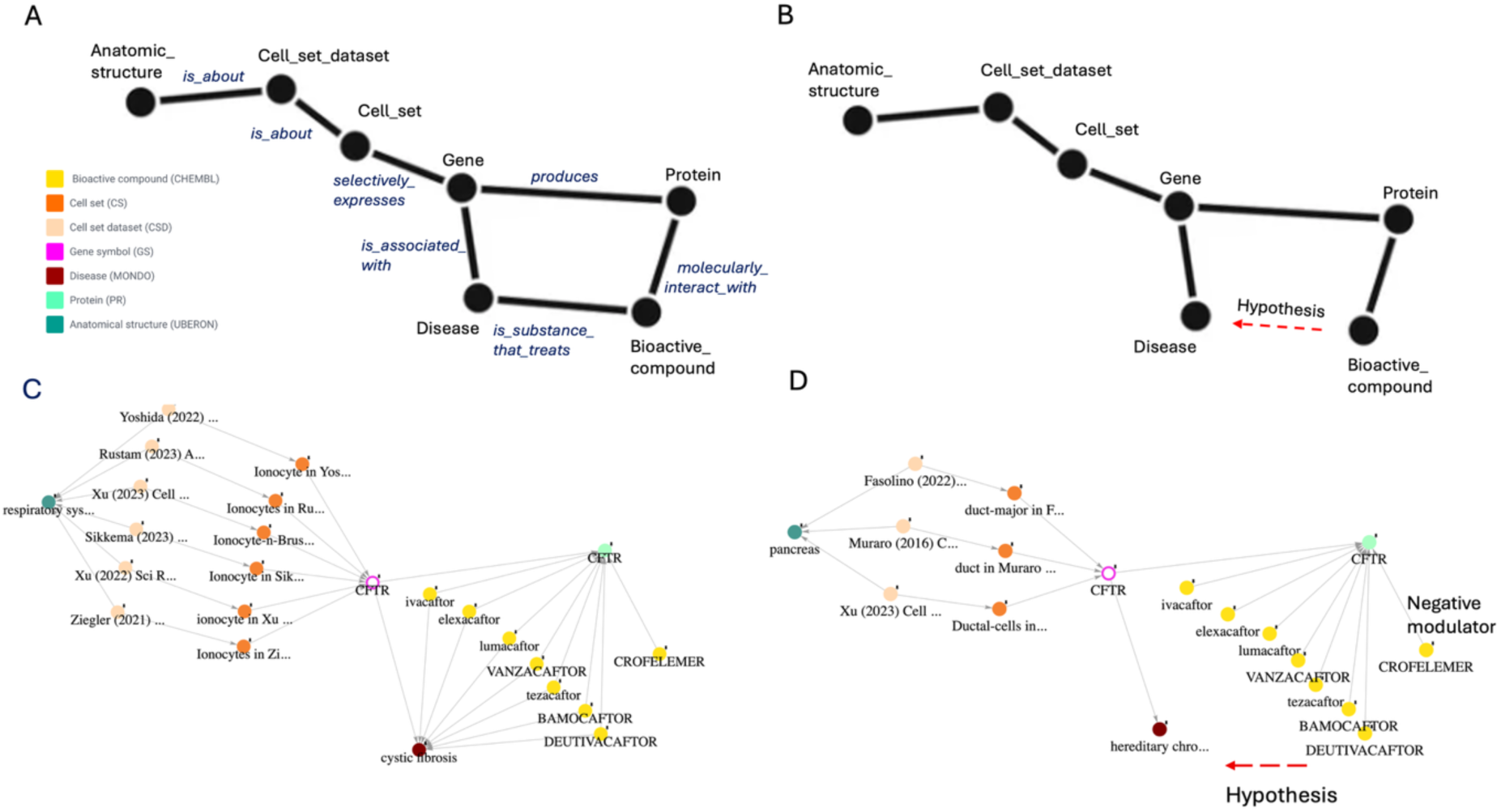
Big Dipper topological structures in NLM-CKN (Use Cases #6 and #7). (**A**) The “Full Big Dipper” topology includes a closed cup composed of a gene that is_associated_with a disease and that produces a protein that molecularly_interacts_with a bioactive compound that is_substance_that_treats_disease. The closed cup is connected to the relevant cell set with the selectively_expresses relation to the gene. (**B**) The “Bottomless Big Dipper” topology includes a cup lacking the bioactive compound is_substance_that_treats disease link. (**C**) Example of a full Big Dipper topology showing that the *CFTR* gene (pink node) is associated with the cystic fibrosis disease (brown node). The *CFTR* protein (cyan node) molecularly interacts with bioactive compounds (yellow nodes) that are used to treat cystic fibrosis. Six independent cell set datasets (tan nodes) identify *CFTR* as selectively expressed in lung ionocyte cell sets (orange nodes). (**D**) Example of a bottomless Big Dipper topology showing that *CFTR* is also associated with hereditary chronic pancreatitis. Three independent cell set datasets identify *CFTR* as selectively expressed in pancreatic ductal cells. The *CFTR* protein molecularly interacts with eight bioactive compounds. Hypothetically, these compounds may have therapeutic potential as drug repurposing candidates for selected patients with *CFTR*-related hereditary chronic pancreatitis, except for crofelemer, which is a negative modulator of the *CFTR* chloride channel.

Figure 6C illustrates an example of the Big Dipper topology and addresses Use Case #6: “Which genetic and cellular factors contribute to cystic fibrosis pathogenesis, and what treatment options are available?” *CFTR* encodes the cystic fibrosis transmembrane conductance regulator, a chloride and bicarbonate ion channel, and is a key causal gene for cystic fibrosis (68,69). In NLM-CKN, *CFTR* is identified as both a marker gene and a binary gene for lung ionocytes across six independent lung datasets. This finding is consistent with previous reports describing the role of ionocytes in maintaining airway ion and fluid homeostasis through *CFTR*-mediated chloride and bicarbonate transport (68,69). The *CFTR*-encoded protein interacts with eight bioactive compounds, seven of which are positive *CFTR* modulators used in approved therapies for cystic fibrosis. In contrast, crofelemer inhibits *CFTR*-mediated chloride transport and is not used to restore *CFTR* function in cystic fibrosis.

To systematically explore the Big Dipper topology, Use Case #9 asks: “List all genes that are selectively expressed in specific cell set(s), are strongly associated with a disease, and encode a protein that interacts with bioactive compound(s) used to treat that disease.” This use case identified 189 genes in NLM-CKN v1.0 that form full Big Dipper topologies, in which all seven node types and their connecting relationships are represented. Supplementary Table S3 lists the gene symbol, category, gene name, and RefSeq summary for these Big Dipper genes. Importantly, these genes encode proteins belonging to several major categories of druggable targets, including G protein-coupled receptors (GPCRs), ion channels, kinases, other receptors, and enzymes.

#### Bottomless Big Dipper

In NLM-CKN, we also identified several Big Dipper-like topologies that lack a connection between bioactive compounds and diseases, as shown in Figure 6B. We refer to these as “bottomless Big Dipper” topologies. Hypothetically, when a gene is associated with a disease and its encoded protein interacts with a bioactive compound, that compound may represent a candidate for drug repurposing to treat the disease.

For example, Figure 6D addresses Use Case #7: “Which genetic and cellular factors contribute to hereditary chronic pancreatitis pathogenesis? What are hypothetical treatments for hereditary chronic pancreatitis?” *CFTR* is not only the primary causal gene for cystic fibrosis but also a recognized genetic causal gene for hereditary chronic pancreatitis (Figure 6D). Dysfunction of *CFTR* impairs epithelial fluid transport in the pancreas, leading to reduced ductal flushing, protein plug formation, ductal obstruction, and recurrent pancreatic inflammation (70). Consistent with this mechanism, three independent pancreas single-cell datasets identified *CFTR* as selectively expressed in pancreatic ductal cells in NLM-CKN. Hypothetically, bioactive compounds that modulate *CFTR* activity and are currently approved for the treatment of cystic fibrosis may also have therapeutic potential for hereditary chronic pancreatitis. This example demonstrates how the bottomless Big Dipper topology can support the identification of potential drug-repurposing opportunities.

It is important to note that not all compounds represented in the bottomless Big Dipper topology are suitable therapeutic candidates for the associated disease. For example, crofelemer is a negative *CFTR* modulator and therefore would not be expected to restore *CFTR* function in hereditary chronic pancreatitis (Figure 6D).

Another example is the *KCNK3* Big Dipper topology shown in Figure 7. *KCNK3*, which encodes the two-pore domain potassium channel TASK-1, is an established disease-associated gene for pulmonary arterial hypertension (PAH) (71). Seven lung datasets identified *KCNK3* as selectively expressed in lung pericytes in NLM-CKN. Pericytes are elongated, contractile cells found wrapped around precapillary arterioles outside the basement membrane (72). NLM-CKN also identifies several bioactive compounds that regulate TASK-1 potassium channel activity. Theoretically, pharmacological modulation of TASK-1 could influence pericyte function and provide a potential therapeutic strategy for PAH. However, several of the compounds linked in NLM-CKN, including halothane, isoflurane, sevoflurane, enflurane, and desflurane, are general anesthetics and are therefore not suitable for the chronic treatment of PAH. Conversely, doxapram inhibits human TASK-1, which would alter channel activity in the opposite direction from that required to improve *KCNK3* function. Thus, no suitable therapeutic candidate was identified in this case.

**Figure 7.**
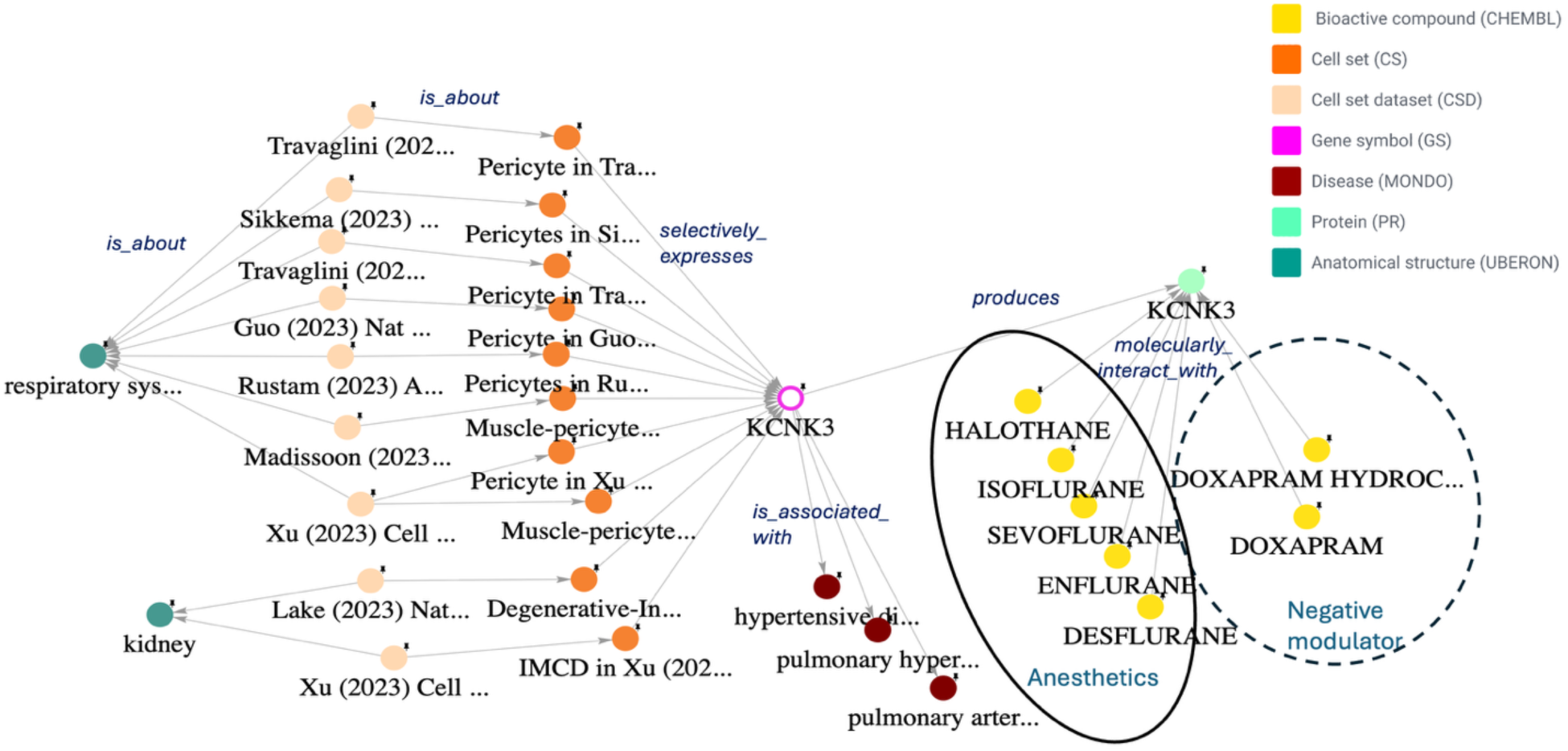
Bottomless Big Dipper topology for *KCNK3* and pulmonary hypertension. *KCNK3*, which encodes the two-pore-domain potassium channel TASK-1, is genetically associated with pulmonary arterial hypertension (PAH) and related pulmonary hypertensive phenotypes (71). *KCNK3* is selectively expressed in lung pericytes and kidney inner medullary collecting duct (IMCD) cells. Several bioactive compounds molecularly interact with the *KCNK3*-encoded protein, including volatile anesthetics (solid oval) and negative modulators (dashed ovalotted circle). However, these volatile anesthetics are not suitable for the chronic treatment of PAH, while doxapram inhibits TASK-1 and therefore acts in the opposite direction from the desired restoration of channel activity. Thus, none of these compounds represents a suitable drug-repurposing candidate for *KCNK3*-associated PAH.

Figure 8 demonstrates a multi-phase exploration of Alzheimer disease using NLM-CKN and addresses Use Case #8: “What genes are strongly associated with Alzheimer disease, and what treatments target these genes?” In the first phase, we identified Alzheimer disease-associated genes represented in NLM-CKN v1.0 (Figure 8A). In the second phase, we attempted to construct Big Dipper topologies for these genes to explore their cellular context and associated therapeutic interventions.

**Figure 8.**
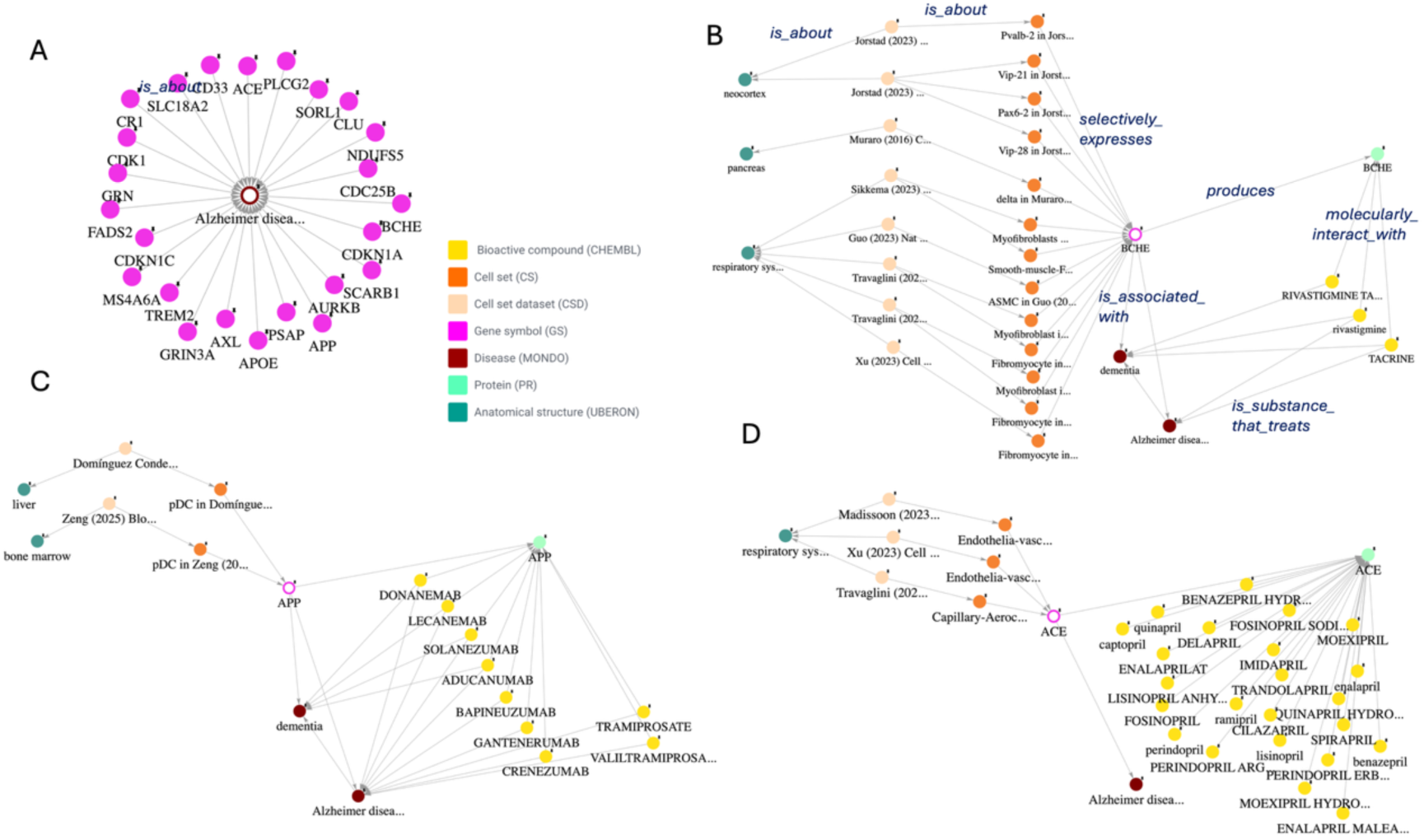
Multi-phase exploration of Alzheimer disease using NLM-CKN (Use Case #8). (**A**) Identification of genes (pink nodes) associated with Alzheimer disease (brown node) represented in NLM-CKN v1.0. (**B**) Bioactive compounds (yellow nodes) targeting the *BCHE*-encoded protein (cyan node) represent established approaches to the symptomatic treatment of Alzheimer disease (brown node), presumably through cell sets (orange nodes) found in the neocortex. Expression of *BCHE* in cell sets in the pancreas and respiratory system could result in adverse treatment side effects. (**C**) Monoclonal antibodies and small molecules targeting *APP*-derived amyloid-β represent newer therapeutic strategies directed at amyloid pathology in Alzheimer disease. *APP* shows selective expression in pDCs in the bone marrow and liver. (**D**) *ACE* inhibitors targeting the *ACE* protein represent a potential therapeutic strategy for Alzheimer disease, but *ACE* is also selectively expressed in endothelial cells in the respiratory system.

Three Alzheimer disease-associated genes – *BCHE, APP*, and *ACE* – form Big Dipper topologies that illustrate traditional, emerging, and potential therapeutic strategies, respectively. NLM-CKN identifies rivastigmine and tacrine as compounds that interact with the *BCHE*-encoded protein and are used to treat Alzheimer disease (Figure 8B), consistent with the traditional cholinesterase-inhibitor approach to the symptomatic treatment of Alzheimer disease (73). NLM-CKN also links *APP* (amyloid precursor protein) to nine monoclonal antibodies and two small molecules used for the treatment of Alzheimer disease (Figure 8C). These relationships are consistent with newer therapeutic strategies directed at *APP*-derived amyloid-β (Aβ): monoclonal antibodies bind specific forms of Aβ and promote their clearance, whereas the small molecules tramiprosate and valiltramiprosate stabilize Aβ monomers and inhibit the formation of neurotoxic Aβ oligomers (74,75). Thus, Figure 8C reflects therapeutic strategies directed at the underlying amyloid pathology of Alzheimer disease. Finally, numerous *ACE* inhibitors have been developed (Figure 8D), and some have been investigated for their potential therapeutic effects in Alzheimer disease; however, the available evidence remains inconclusive, and *ACE* inhibitors are not currently approved specifically for the treatment of Alzheimer disease (76).

Use Case #10 asks: “List all genes that are selectively expressed in specific cell set(s), are strongly associated with a disease, and encode a protein that interacts with bioactive compound(s).” This use case identified 442 genes in NLM-CKN v1.0 that form bottomless Big Dipper topologies, in which the link between the disease and the bioactive compound is absent. Supplementary Table S4 lists the gene symbol, category, gene name, and RefSeq summary for these genes. In addition, the workflow builder provides a workflow for exploring individual Big Dipper candidate genes by generating the corresponding topology for each gene.

### Classes of Big Dipper topologies

Since NLM-CKN is a cell-focused knowledge network that integrates single-cell expression data with knowledge of disease mechanisms and therapeutic interventions, it enables investigation of cellular contributions to disease pathogenesis and the cellular context of therapeutic interventions. Because of the complexity of gene-disease relationships, different classes of Big Dipper topologies require distinct biological interpretations.

#### Big Dipper topology for exploring cellular contributions to disease mechanisms

Class 1 represents Big Dipper topologies in which the selectively expressed gene is genetically associated with the disease, and disease-manifesting cells are among the cells expressing the disease-associated gene. In these cases, the Big Dipper provides meaningful evidence for identifying potential cellular contributors to disease pathogenesis. However, there are incomplete Class 1 topologies in which disease-manifesting cells are not among the identified cells expressing the disease-associated gene, likely reflecting incomplete single-cell data or broader gene expression not captured in NLM-CKN v1.0 and therefore requiring cautious interpretation.

Figure 9A presents an example of a Class 1 topology and addresses Use Case #11, which asks: “What is the contribution of retinal pigment epithelial (RPE) cells to *RPE65*-related Leber congenital amaurosis?” In this example, *RPE65* is a causal gene for *RPE65*-related retinitis pigmentosa and Leber congenital amaurosis. Loss of RPE65 reduces the production of 11-cis-retinal required by photoreceptors, ultimately leading to visual dysfunction and retinal degeneration (77). Voretigene neparvovec is a gene therapy that treats these conditions by restoring RPE65 protein function (78). NLM-CKN identifies *RPE65* as a marker gene for RPE cells, and its expression is largely restricted to RPE cells. Thus, RPE cells are key cellular contributors to *RPE65*-related disease pathogenesis.

**Figure 9.**
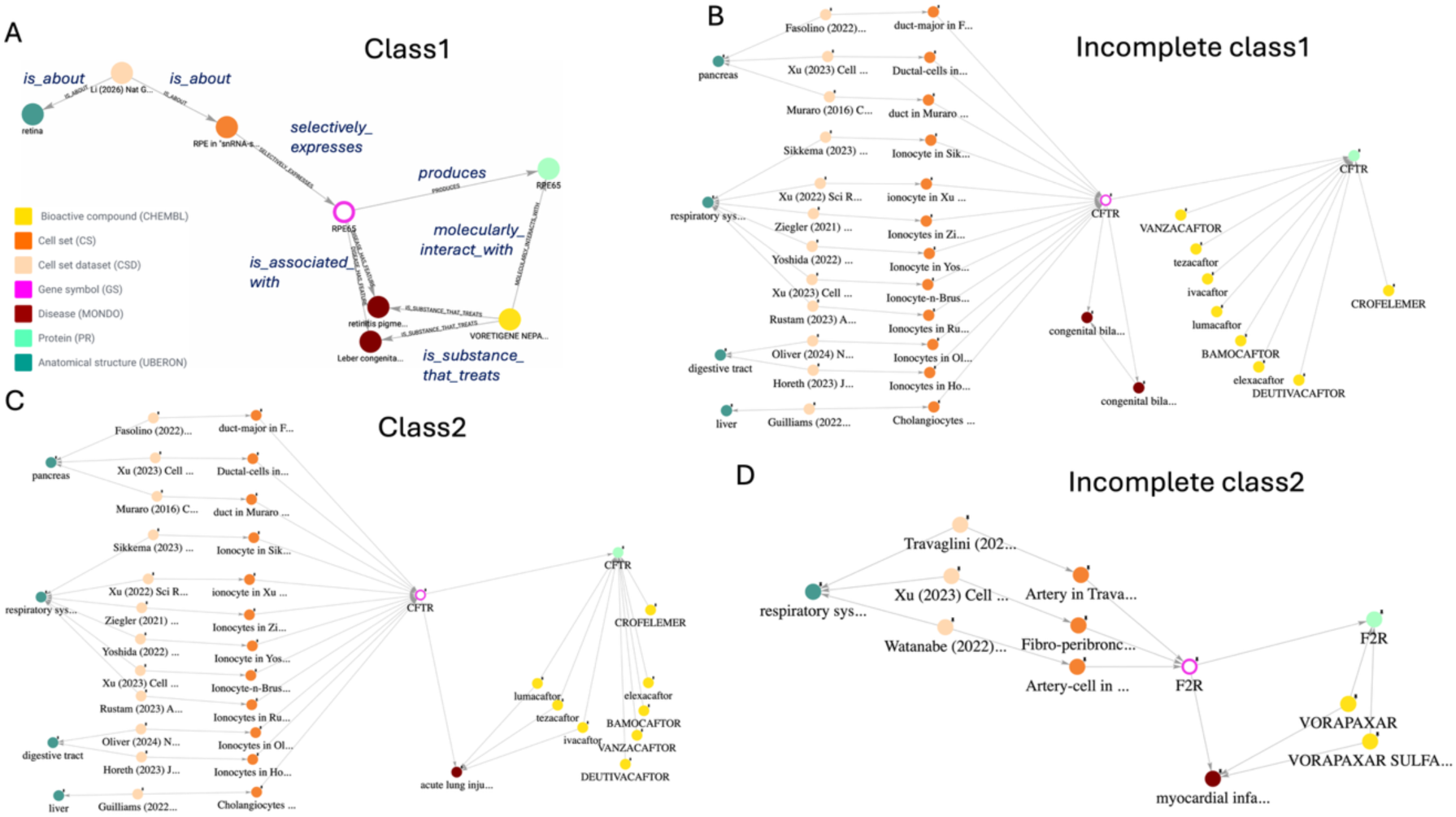
Examples of the two classes of Big Dipper topologies. (**A**) Class 1: *RPE65* is a causal gene for *RPE65*-related Leber congenital amaurosis (brown nodes) and is selectively expressed in retinal pigment epithelial (RPE) cells, supporting RPE cells as a cellular driver of disease pathogenesis. (**B**) Incomplete Class 1: *CFTR* is a causal gene for congenital bilateral absence of the vas deferens (brown nodes). However, because vas deferens data are not included in NLM-CKN v1.0, the topology cannot be used to assess the cellular contribution to this disease. (**C**) Class 2: *CFTR* is associated with acute lung injury and is selectively expressed in multiple cell types, including lung ionocytes. Therefore, compounds targeting *CFTR* may affect multiple *CFTR*-expressing cell types across different tissues, including lung and digestive tract ionocytes and liver and pancreatic ductal cells. (**D**) Incomplete Class 2: *F2R* is associated with myocardial infarction, but selective *F2R* expression in NLM-CKN v1.0 is identified only in cell sets from the respiratory system. This discrepancy may reflect incomplete data coverage; therefore, the cellular and tissue context of *F2R*-targeted therapies should be interpreted cautiously.

The *CFTR* examples in Figures 6B and 6D also represent Class 1 topologies. *CFTR* is a causal gene for diseases including cystic fibrosis, hereditary chronic pancreatitis, and congenital bilateral absence of the vas deferens (79). Consistent with this, the phenotypes associated with *CFTR* mutations manifest in the lung, pancreas, and vas deferens. *CFTR* is a marker gene for lung ionocytes and pancreatic ductal cells, aligning the cellular locations of disease with the cell sets that express the causal disease gene. Thus, lung ionocytes and pancreatic ductal cells represent key cellular contributors to the pathogenesis of cystic fibrosis and hereditary chronic pancreatitis, respectively.

Notably, *CFTR* expression in vas deferens cells is not represented in the current KG because the vas deferens is not included among the 10 anatomical systems currently covered by NLM-CKN v1.0. Consequently, the Big Dipper involving congenital bilateral absence of the vas deferens represents an incomplete Class 1 topology, in which the known disease-manifesting tissue is not represented among the identified *CFTR*-expressing cells (Figure 9B). However, *CFTR* expression in the vas deferens has been reported in the literature (79), suggesting that *CFTR*-expressing cell types in the vas deferens may be the cellular contributors to congenital bilateral absence of the vas deferens. This example demonstrates the importance of considering KG coverage when interpreting Big Dippers.

The Big Dipper in Figure 8C is another example of a Class 1 topology. *APP* has an established genetic role in Alzheimer disease. Previous studies have reported that *APP* is a neuron-specific protein under basal and neuroinflammatory conditions (80). However, NLM-CKN v1.0 identifies *APP* as a selectively expressed gene only in plasmacytoid dendritic (pDC) cell sets from the healthy liver and bone marrow. Therefore, additional expression data are needed to interpret the cellular contributions of neocortical cell types to Alzheimer disease pathogenesis.

#### Big Dipper topology for exploring the cellular context of therapeutic intervention

Class 2 represents Big Dipper topologies in which the selectively expressed gene is not genetically associated with the disease but is a therapeutic target for the symptomatic treatment of the disease, and disease-relevant cell types are among the cell types expressing the therapeutic target gene. In these cases, the Big Dipper provides useful information about the cellular and tissue contexts in which the therapeutic target may act and provides insights into potential off-target effects. However, in incomplete Class 2 topologies, disease-relevant cell types may not be among those expressing the therapeutic target gene. This discrepancy may reflect insufficient expression data or incomplete representation of relevant cell types in NLM-CKN v1.0. Therefore, the inferred cellular and tissue contexts of the therapeutic target should be interpreted cautiously.

Figure 9C addresses Use Case #12, which asks: “What is the cellular context when using CFTR-targeted therapy to treat acute lung injury?” *CFTR* is a disease-associated gene in acute lung injury. Three compounds represented in NLM-CKN that improve *CFTR* channel activity and enhance alveolar liquid clearance are approved to treat the symptoms of acute lung injury. NLM-CKN identifies *CFTR* as selectively expressed in ionocytes, ductal cells, and cholangiocytes across the respiratory system, digestive tract, pancreas, and liver, providing cellular and tissue contexts for *CFTR*-targeted therapeutic interventions. Therefore, Figure 9C represents a Class 2 Big Dipper topology.

Figure 8B is another example of a Class 2 topology. *BCHE* is associated with Alzheimer disease and encodes butyrylcholinesterase (BChE). The compounds shown in NLM-CKN inhibit cholinesterase activity, increasing the availability of acetylcholine and thereby improving cognitive symptoms in Alzheimer disease (73). Several GABAergic neuronal cell sets in the neocortex selectively express *BCHE*, providing a disease-relevant cellular context for the symptomatic treatment of Alzheimer disease. In addition to these neuronal populations, *BCHE* is selectively expressed in cell sets in the respiratory system and pancreas, providing broader cellular and tissue contexts for *BCHE*-targeted therapies.

Figure 9D is an example of an incomplete Class 2 topology. *F2R* encodes PAR-1, a thrombin receptor that mediates platelet activation. The PAR-1 antagonist vorapaxar reduces recurrent cardiovascular events in patients with prior myocardial infarction (81). This therapy is captured in NLM-CKN, but selective *F2R* expression in NLM-CKN v1.0 is identified only in cell sets from the respiratory system. This discrepancy may reflect incomplete data coverage; therefore, the cellular and tissue contexts of *F2R*-targeted therapies should be interpreted cautiously.

As mentioned above, *ACE* is associated with Alzheimer disease, and brain-penetrating *ACE* inhibitors have been investigated for their potential effects on cognitive decline in patients with Alzheimer disease (76). NLM-CKN detects selective *ACE* expression in endothelial cell sets in the respiratory system; however, disease-relevant brain cell types expressing *ACE* are not represented in NLM-CKN v1.0. Therefore, Figure 8D represents an example of an incomplete Class 2 topology. The cellular context of *ACE*-targeted therapies should be carefully evaluated when considering their potential application to Alzheimer disease.

Additional examples illustrating the different Big Dipper topology classes are provided in Supplementary Figures S5-S9.

## Discussions

### Marker genes and binary genes in NLM-CKN

NLM-CKN v1.0 contains 5,139 genes designated as selectively expressed in cell sets for a given anatomical structure, which are components of biomarker combinations and binary gene sets nodes. Genes included in biomarker combinations are referred to as marker genes, whereas genes included in binary gene sets are referred to as binary genes. Both marker genes and binary genes are identified by using the validated computational pipeline NS-Forest and supported by experimental evidence and quantitative statistical confidence, including F-beta scores and binary scores.

NLM-CKN marker genes differ from many classical cell-type markers established before the advent of single-cell technologies, when cell types were generally classified using morphology and/or protein markers assessed by immunohistochemistry or flow cytometry at coarser levels of granularity. By representing marker genes that are systematically derived from high-content single-cell data using computational algorithms, NLM-CKN provides cell-set-specific molecular signatures at finer resolution, enabling several downstream applications.

In particular, the biomarker combinations generated by the NS-Forest pipeline identify necessary and sufficient combinations of marker genes for distinguishing individual cell sets. Collectively, these combinations constitute a minimal, non-redundant transcript panel that can facilitate the design of targeted spatial transcriptomics probes. For example, the 115 biomarker combination gene panel in Supplementary Table S2 could be used for optimal spatial resolution of cell-types in the human lung. Marker genes can also be used to refine Cell Ontology term definitions and naming conventions. Indeed, the Cell Ontology is currently incorporating NS-Forest markers, along with their associated F-beta scores, into cell type descriptions for cell types found in single-cell transcriptomics datasets. Finally, marker genes can support cross-dataset mapping and cell annotation. They can define reduced but informative feature spaces, enabling more efficient mapping of cell sets across datasets. This concept has been applied in the development of FR-Match, one of the most reliable methods for matching cell sets across datasets (49,60).

Because NS-Forest optimizes for a minimal combination sufficient for cell set classification, each cell set has a small number of marker genes, typically between two and three. While this minimal representation is advantageous for classification, a larger set of selectively expressed genes can benefit applications that require a more comprehensive representation of gene expression, such as characterization of cell type function and identity. Therefore, NLM-CKN v1.0 also includes the top 10 binary genes for each cell set. These additional genes retain cell set-selective expression features while providing complementary molecular information for characterizing cell phenotypes and exploring topological relationships within NLM-CKN.

### Grounding Cell Ontology terms in experimental evidence

A key feature of NLM-CKN is the anchoring of Cell Ontology (CL) cell type terms in experimentally derived cell-set evidence. Because CL is hierarchically organized, mapping experimental cell sets to CL terms places them within a standardized cell-type classification framework and enables users to query and aggregate cell sets at different levels of granularity. For example, Figure 4 demonstrates how four dendritic cell sets from the HLCA dataset can be organized according to the corresponding dendritic cell hierarchies in CL.

Conversely, experimentally derived cell sets can provide evidence for refinement of the Cell Ontology. Cell populations that cannot be adequately mapped to existing CL terms may reveal gaps, ambiguities, or insufficiently resolved definitions, while reproducible cell phenotypes observed across independent datasets can provide evidence for improved cell-type characterization. For example, Figure 3 shows that nasal club cells and non-nasal club cells are distinguished as separate experimental cell sets, although both currently map to the same CL term, “club cell”. If additional datasets consistently support a distinct nasal club cell phenotype, the Cell Ontology could consider representing club cell subtypes at a more granular level. In addition, Figure 6C shows that ionocyte cell sets from six independent datasets consistently identify *CFTR* as a marker gene or binary gene, providing reproducible experimental evidence supporting the characterization of pulmonary ionocytes.

On closer examination of Figure 4, DC1 and DC2 are two distinct experimental cell sets that map to conventional dendritic cell and CD1c-positive myeloid dendritic cell, respectively. However, in the current Cell Ontology hierarchy, conventional dendritic cell is the parent of myeloid dendritic cell, which is the parent of CD1c-positive myeloid dendritic cell. Given the experimental evidence that DC1 and DC2 represent distinct cell populations, the relationships in CL among conventional dendritic cell, myeloid dendritic cell, and CD1c-positive myeloid dendritic cell, as well as their relationships to the common ancestor dendritic cell, may warrant adjustment.

### Recurrent semantic topologies

The “Big Dipper” topology links diseases and therapeutic interventions (the bucket of the Big Dipper) with single-cell knowledge (the handle of the Big Dipper). These recurrent semantic topologies form the basis for several types of hypotheses.

First, as shown in Figure 6B, the “bottomless” Big Dipper topology represents drug-repurposing hypotheses. When a gene is strongly associated with a disease and its encoded protein molecularly interacts with a compound, that compound may be considered a potential candidate for treating the disease. However, as illustrated in Figure 7, such hypotheses must be evaluated carefully because some compounds may exert effects that are contraindicated for treating the disease or may cause undesirable side effects. In addition, the type, direction, and strength of gene-disease associations can affect the confidence of drug-repurposing predictions. For example, *ACE* has a complex association with Alzheimer disease. *ACE* plays multiple, potentially competing roles in the brain and influences inflammation, vascular tone, and the clearance of amyloid-beta (Aβ). These complex gene-disease mechanisms complicate the prediction of therapeutic effects of *ACE*-targeting compounds. In this regard, we are exploring strategies to prioritize drug-repurposing candidates by considering such complexities in gene-disease and drug-target relationships (82).

The bucket of the Big Dipper has been modeled in several existing biomedical KGs. The novel contribution of NLM-CKN is the handle of the Big Dipper and its connection to the bucket, which provides evidence for cellular contributions to disease pathogenesis. In Class 1 Big Dipper topologies, a gene genetically associated with a disease is selectively expressed in a particular cell set and its corresponding cell type. Linking disease genetics to cell set-specific gene expression can therefore identify candidate cell populations that contribute directly to disease pathogenesis. For example, *CFTR* is genetically associated with cystic fibrosis and selectively expressed in lung ionocytes, which exhibit high channel activity and contribute to the regulation of ion/fluid homeostasis in the airway, supporting their role as cellular contributors to cystic fibrosis pathogenesis (68,69).

In Class 2 Big Dipper topologies, a gene is merely associated with a disease and may serve as a therapeutic target to alleviate disease *symptoms*. We cannot infer cellular contributions to disease development based on this class of Big Dipper topologies; however, we can predict the cell types in which the drug may act to relieve symptoms and the potential off-target effects of the treatment. For example, when using the compounds in Figure 9C to treat acute lung disease, these compounds could also affect cells in the liver, pancreas, and digestive tract.

As noted for incomplete Class 1 and 2 topologies, incomplete cellular gene-expression information in NLM-CKN v1.0 limits its ability to provide a comprehensive cellular context for disease-associated genes. Two coverage limitations contribute to this issue. First, NLM-CKN v1.0 includes single-cell datasets from 10 anatomical systems; therefore, cellular expression profiles for genes in other organs and tissues are not represented. Second, v1.0 primarily models selectively expressed genes rather than comprehensive gene expression within each cell set. Specifically, Cell_Set-Selectively_Expresses-Gene assertions are limited to the top 10 binary genes for each cell set, although a cell population typically expresses thousands of genes. Consequently, some disease-associated genes may be absent from the cellular expression layer of the current KG. Thus, using disease-associated genes to identify cell sets in which a disease may manifest can provide only a partial picture when relevant gene-expression relationships are not represented. On the other hand, the exquisite cell type selectivity of these genes could reflect useful therapeutic properties.

### Future Work

NLM-CKN v1.0 covers 10 anatomical systems. In future versions, we plan to expand NLM-CKN’s coverage to organs and tissues for which atlas data are published for the Human Cell Atlas (HCA), including adipose tissue, male and female reproductive systems, breast, musculoskeletal tissues, and oral and craniofacial tissues.

Additionally, future versions of NLM-CKN will represent all expressed genes for each cell set, rather than focusing primarily on selectively expressed genes. This expanded representation will provide a more comprehensive molecular view of cell phenotype and enable identification of biological processes and pathways operating in each cell set. Importantly, it will also provide more complete expression evidence for identifying disease-manifesting cell populations by linking disease-associated genes to the cell types in which they are expressed, thereby improving the characterization of disease mechanisms and therapeutic targets.

In NLM-CKN v1.0, cell sets from 10 reference datasets were manually mapped to Cell Ontology (CL) terms. Manual cell set-to-cell type mapping relies on domain expertise and is time-consuming and error-prone. Therefore, once cell sets from reference datasets have been mapped to cell types, we plan to use a computational cell-set matching framework to minimize manual mapping and support the incremental growth of the knowledgebase. This framework incorporates FR-Match and additional computational cell-set matching methods evaluated in our benchmarking study (60). Cell sets from new query datasets will be computationally matched against those in established reference datasets. Exact matches can inherit the CL associations of the corresponding reference cell sets, whereas related but non-exact matches should be evaluated to determine whether they represent more specific or broader cell types. Query cell sets that do not match but have sufficient evidence to support a distinct cell phenotype can be incorporated as new reference cell sets and proposals for new cell types in CL. As a demonstration in NLM-CKN v1.0, CellRef cell sets were computationally matched against the HLCA reference dataset. Unmatched or partially matched cell sets can be evaluated for incorporation into the reference, enabling NLM-CKN to expand as new datasets become available.

Future versions of NLM-CKN will begin incorporating single-cell datasets from diseased tissues. Differential gene-expression analyses will allow the KG to represent disease-related changes at cell-type resolution and facilitate investigation of cell-type-specific molecular alterations associated with disease pathogenesis and therapeutic intervention.

As single-cell technologies expand to single-cell multiomics (13–15) and spatial transcriptomics (17,18,33), NLM-CKN will extend beyond gene-expression-based cell phenotypes to capture the regulatory mechanisms underlying cell phenotypes and cell-cell interactions. A future version of NLM-CKN will represent spatial associations among cell sets, cell-cell communication, and regulatory relationships linking chromatin accessibility and regulatory elements with transcription factor activity, target gene expression, biological pathways, and overall cell phenotypes.

In addition to the cell characteristics derived from the validated computational pipelines currently used in NLM-CKN, complementary knowledge will also be extracted from peer-reviewed primary literature using structured Large Language Models/Natural Language Processing (LLM/NLP) approaches. Our group developed CellLink, a reusable benchmark for developing and evaluating cell-population named entity recognition and ontology linking systems for ontology-aware literature curation (83). Building on this foundation, future versions of NLM-CKN will incorporate LLM/NLP-generated knowledge triples to capture additional cell phenotype knowledge reported in the literature.

At the infrastructure level, future development will pursue two complementary strategies to connect NLM-CKN with broader biomedical knowledge resources. Key resources will be directly integrated through semantic and schema harmonization, including alignment of entities, relationships, identifiers, and ontologies. In addition, federated integration will enable cross-resource queries while allowing external knowledge to remain within its source databases. Together, these approaches will establish a comprehensive, interoperable biomedical knowledge network.

To enable intuitive access to NLM-CKN and explore our use cases, we are developing an AI agent that allows users to submit biomedical questions in natural language. An LLM host, equipped with knowledge of the NLM-CKN schema, interprets each question and either maps it to a predefined query or dynamically generates an ArangoDB Query Language (AQL) query tailored to the user’s request. The query is then executed by a Model Context Protocol (MCP) server, which runs the canned or generated AQL query against NLM-CKN and returns the relevant subgraph. Finally, an LLM agent synthesizes the retrieved subgraph into a coherent natural language response, accompanied by citations to the underlying data sources.

To make NLM-CKN AI-ready, future versions will provide complementary representations of KG content for downstream AI applications. First, SSS assertions will be textualized by converting structured assertions into controlled natural-language statements (e.g., “ionocytes selectively express *CFTR*”), making KG knowledge directly consumable by LLMs. Second, SSS assertions will be serialized into structured representations that explicitly encode entities, relations, ontology identifiers, and biological context, providing machine-readable inputs that can be tokenized for model training. Third, embeddings will be generated for entities, relations, and biologically meaningful subgraphs to support semantic retrieval, similarity search, link prediction, and retrieval-augmented generation. Together, these representations will make NLM-CKN knowledge more accessible to AI models.

## Data availability

All processed data including filtered `.h5ad` files are freely available for academic use and can be downloaded from S3 bucket (https://nlm-ckn.nlm.nih.gov/V1.0) and Zenodo (https://doi.org/10.5281/zenodo.23407774). User feedback, issue reporting, and code access are supported through GitHub (https://github.com/NIH-NLM/nlm-ckn) and Zenodo.

## Acknowledgements

We thank Chris Mungall, Belinda Seto, Vidhya Venkateswaran, Vik Becker, Kellen Doggett, Matthew Nho, Kristen Oguno, Kristine Pashin for useful discussion. This research was supported in part by the Intramural Research Program of the National Institutes of Health (NIH) (ZIA-LM202401, ZIA-LM202501). The contributions of the NIH authors are considered Works of the United States Government. The findings and conclusions presented in this paper are those of the authors and do not necessarily reflect the views of the NIH or the U.S. Department of Health and Human Services.

## Author contributions

Bingfang Xu (Conceptualization [lead], Data curation [equal], Validation [equal], Methodology [equal], Project administration [equal], Writing – original draft [lead]), Anne Deslattes-Mays (Conceptualization [equal], Data curation [equal], Validation [equal], Methodology [equal], Software [equal], Project administration [lead], Writing – original draft [equal]), Matthew Diller (Conceptualization [lead], Validation [equal], Methodology [equal], Writing – original draft [equal]), Raymond LeClair (Conceptualization [equal], Data curation [equal], Methodology [equal], Software [equal], Writing – original draft [equal]), Ajith V. Pankajam (Validation [equal], Methodology [equal], Software [lead]), William Spear (Validation [equal], Methodology [equal], Software [equal], Writing – original draft [equal]), David Osumi-Sutherland (Conceptualization [equal], Validation [equal],), Kumar Arvind (Validation [equal]), Sreejato Chatterjee (Validation [equal]), Alexander Diehl (Validation [equal]), Md Mezbahul Islam (Validation [equal]), Conor Kelly (Validation [equal]), Rana Khalil (Validation [equal]), Juhi Khare (Vizualization [equal]), Ricardo Lopez Candelaria (Validation [equal]), Beverly Peng (Methodology [equal], Validation [equal]), Latoya McDonald (Validation [equal]), Daniel Pintard (Validation [equal]), Noam Rotenberg (Data curation [equal]), David Zweibaum (Validation [equal]), Yun Zhang (Conceptualization [lead], Data curation [lead], Validation [equal], Methodology [lead], Supervision [equal], Writing – review & editing [equal], Funding acquisition [equal]), Richard H. Scheuermann (Conceptualization [lead], Data curation [lead], Validation [equal], Supervision [lead], Writing – review & editing [lead], Funding acquisition [lead]))

## Author notes

This is an Open Access article distributed under the terms of the Creative Commons Attribution License (https://creativecommons.org/licenses/by/4.0/), which permits unrestricted reuse, distribution, and reproduction in any medium, provided the original work is properly cited.

## References

1. Buniello, A., MacArthur, J.A.L., Cerezo, M., Harris, L.W., Hayhurst, J., Malangone, C., McMahon, A., Morales, J., Mountjoy, E., Sollis, E. et al. (2019) The NHGRI-EBI GWAS Catalog of published genome-wide association studies, targeted arrays and summary statistics 2019. Nucleic Acids Res, 47, D1005–D1012.

2. Claussnitzer, M., Cho, J.H., Collins, R., Cox, N.J., Dermitzakis, E.T., Hurles, M.E., Kathiresan, S., Kenny, E.E., Lindgren, C.M., MacArthur, D.G. et al. (2020) A brief history of human disease genetics. Nature, 577, 179–189.

3. Tam, V., Patel, N., Turcotte, M., Bosse, Y., Pare, G. and Meyre, D. (2019) Benefits and limitations of genome-wide association studies. Nat Rev Genet, 20, 467–484.

4. Visscher, P.M., Wray, N.R., Zhang, Q., Sklar, P., McCarthy, M.I., Brown, M.A. and Yang, J. (2017) 10 Years of GWAS Discovery: Biology, Function, and Translation. Am J Hum Genet, 101, 5–22.

5. Tang, F., Barbacioru, C., Wang, Y., Nordman, E., Lee, C., Xu, N., Wang, X., Bodeau, J., Tuch, B.B., Siddiqui, A. et al. (2009) mRNA-Seq whole-transcriptome analysis of a single cell. Nat Methods, 6, 377–382.

6. Islam, S., Kjallquist, U., Moliner, A., Zajac, P., Fan, J.B., Lonnerberg, P. and Linnarsson, S. (2011) Characterization of the single-cell transcriptional landscape by highly multiplex RNA-seq. Genome Res, 21, 1160–1167.

7. Hashimshony, T., Wagner, F., Sher, N. and Yanai, I. (2012) CEL-Seq: single-cell RNA-Seq by multiplexed linear amplification. Cell Rep, 2, 666–673.

8. Picelli, S., Bjorklund, A.K., Faridani, O.R., Sagasser, S., Winberg, G. and Sandberg, R. (2013) Smart-seq2 for sensitive full-length transcriptome profiling in single cells. Nat Methods, 10, 1096–1098.

9. Krishnaswami, S.R., Grindberg, R.V., Novotny, M., Venepally, P., Lacar, B., Bhutani, K., Linker, S.B., Pham, S., Erwin, J.A., Miller, J.A. et al. (2016) Using single nuclei for RNA-seq to capture the transcriptome of postmortem neurons. Nat Protoc, 11, 499–524.

10. Macosko, E.Z., Basu, A., Satija, R., Nemesh, J., Shekhar, K., Goldman, M., Tirosh, I., Bialas, A.R., Kamitaki, N., Martersteck, E.M. et al. (2015) Highly Parallel Genome-wide Expression Profiling of Individual Cells Using Nanoliter Droplets. Cell, 161, 1202–1214.

11. Zheng, G.X., Terry, J.M., Belgrader, P., Ryvkin, P., Bent, Z.W., Wilson, R., Ziraldo, S.B., Wheeler, T.D., McDermott, G.P., Zhu, J. et al. (2017) Massively parallel digital transcriptional profiling of single cells. Nat Commun, 8, 14049.

12. Svensson, V., Vento-Tormo, R. and Teichmann, S.A. (2018) Exponential scaling of single-cell RNA-seq in the past decade. Nat Protoc, 13, 599–604.

13. Buenrostro, J.D., Wu, B., Litzenburger, U.M., Ruff, D., Gonzales, M.L., Snyder, M.P., Chang, H.Y. and Greenleaf, W.J. (2015) Single-cell chromatin accessibility reveals principles of regulatory variation. Nature, 523, 486–490.

14. Stoeckius, M., Hafemeister, C., Stephenson, W., Houck-Loomis, B., Chattopadhyay, P.K., Swerdlow, H., Satija, R. and Smibert, P. (2017) Simultaneous epitope and transcriptome measurement in single cells. Nat Methods, 14, 865–868.

15. Cao, J., Cusanovich, D.A., Ramani, V., Aghamirzaie, D., Pliner, H.A., Hill, A.J., Daza, R.M., McFaline-Figueroa, J.L., Packer, J.S., Christiansen, L. et al. (2018) Joint profiling of chromatin accessibility and gene expression in thousands of single cells. Science, 361, 1380–1385.

16. Moses, L. and Pachter, L. (2022) Museum of spatial transcriptomics. Nat Methods, 19, 534–546.

17. Chen, K.H., Boettiger, A.N., Moffitt, J.R., Wang, S. and Zhuang, X. (2015) RNA imaging. Spatially resolved, highly multiplexed RNA profiling in single cells. Science, 348, aaa6090.

18. Stahl, P.L., Salmen, F., Vickovic, S., Lundmark, A., Navarro, J.F., Magnusson, J., Giacomello, S., Asp, M., Westholm, J.O., Huss, M. et al. (2016) Visualization and analysis of gene expression in tissue sections by spatial transcriptomics. Science, 353, 78–82.

19. Fullgrabe, A., George, N., Green, M., Nejad, P., Aronow, B., Fexova, S.K., Fischer, C., Freeberg, M.A., Huerta, L., Morrison, N. et al. (2020) Guidelines for reporting single-cell RNA-seq experiments. Nat Biotechnol, 38, 1384–1386.

20. Rood, J.E., Maartens, A., Hupalowska, A., Teichmann, S.A. and Regev, A. (2022) Impact of the Human Cell Atlas on medicine. Nat Med, 28, 2486–2496.

21. Borner, K., Teichmann, S.A., Quardokus, E.M., Gee, J.C., Browne, K., Osumi-Sutherland, D., Herr, B.W., 2nd, Bueckle, A., Paul, H., Haniffa, M. et al. (2021) Anatomical structures, cell types and biomarkers of the Human Reference Atlas. Nat Cell Biol, 23, 1117–1128.

22. Jain, S., Pei, L., Spraggins, J.M., Angelo, M., Carson, J.P., Gehlenborg, N., Ginty, F., Goncalves, J.P., Hagood, J.S., Hickey, J.W. et al. (2023) Advances and prospects for the Human BioMolecular Atlas Program (HuBMAP). Nat Cell Biol, 25, 1089–1100.

23. Program, C.Z.I.C.S., Abdulla, S., Aevermann, B., Assis, P., Badajoz, S., Bell, S.M., Bezzi, E., Cakir, B., Chaffer, J., Chambers, S. et al. (2025) CZ CELLxGENE Discover: a single-cell data platform for scalable exploration, analysis and modeling of aggregated data. Nucleic Acids Res, 53, D886–D900.

24. Moreno, P., Fexova, S., George, N., Manning, J.R., Miao, Z., Mohammed, S., Munoz-Pomer, A., Fullgrabe, A., Bi, Y., Bush, N. et al. (2022) Expression Atlas update: gene and protein expression in multiple species. Nucleic Acids Res, 50, D129–D140.

25. Clough, E., Barrett, T., Wilhite, S.E., Ledoux, P., Evangelista, C., Kim, I.F., Tomashevsky, M., Marshall, K.A., Phillippy, K.H., Sherman, P.M. et al. (2024) NCBI GEO: archive for gene expression and epigenomics data sets: 23-year update. Nucleic Acids Res, 52, D138–D144.

26. Athar, A., Fullgrabe, A., George, N., Iqbal, H., Huerta, L., Ali, A., Snow, C., Fonseca, N.A., Petryszak, R., Papatheodorou, I., et al. (2019) ArrayExpress update - from bulk to single-cell expression data. Nucleic Acids Res, 47, D711–D715.

27. Network, B.I.C.C. (2021) A multimodal cell census and atlas of the mammalian primary motor cortex. Nature, 598, 86–102.

28. Ament, S.A., Adkins, R.S., Carter, R., Chrysostomou, E., Colantuoni, C., Crabtree, J., Creasy, H.H., Degatano, K., Felix, V., Gandt, P. et al. (2023) The Neuroscience Multi-Omic Archive: a BRAIN Initiative resource for single-cell transcriptomic and epigenomic data from the mammalian brain. Nucleic Acids Res, 51, D1075–D1085.

29. Sikkema, L., Ramirez-Suastegui, C., Strobl, D.C., Gillett, T.E., Zappia, L., Madissoon, E., Markov, N.S., Zaragosi, L.E., Ji, Y., Ansari, M. et al. (2023) An integrated cell atlas of the lung in health and disease. Nat Med, 29, 1563–1577.

30. Guo, M., Morley, M.P., Jiang, C., Wu, Y., Li, G., Du, Y., Zhao, S., Wagner, A., Cakar, A.C., Kouril, M. et al. (2023) Guided construction of single cell reference for human and mouse lung. Nat Commun, 14, 4566.

31. Hansen, J., Sealfon, R., Menon, R., Eadon, M.T., Lake, B.B., Steck, B., Anjani, K., Parikh, S., Sigdel, T.K., Zhang, G. et al. (2022) A reference tissue atlas for the human kidney. Sci Adv, 8, eabn4965.

32. Regev, A., Teichmann, S.A., Lander, E.S., Amit, I., Benoist, C., Birney, E., Bodenmiller, B., Campbell, P., Carninci, P., Clatworthy, M. et al. (2017) The Human Cell Atlas. Elife, 6.

33. Stuart, T. and Satija, R. (2019) Integrative single-cell analysis. Nat Rev Genet, 20, 257–272.

34. Morris, J.H., Soman, K., Akbas, R.E., Zhou, X., Smith, B., Meng, E.C., Huang, C.C., Cerono, G., Schenk, G., Rizk-Jackson, A. et al. (2023) The scalable precision medicine open knowledge engine (SPOKE): a massive knowledge graph of biomedical information. Bioinformatics, 39.

35. Wood, E.C., Glen, A.K., Kvarfordt, L.G., Womack, F., Acevedo, L., Yoon, T.S., Ma, C., Flores, V., Sinha, M., Chodpathumwan, Y. et al. (2022) RTX-KG2: a system for building a semantically standardized knowledge graph for translational biomedicine. BMC Bioinformatics, 23, 400.

36. Morton, K., Wang, P., Bizon, C., Cox, S., Balhoff, J., Kebede, Y., Fecho, K. and Tropsha, A. (2019) ROBOKOP: an abstraction layer and user interface for knowledge graphs to support question answering. Bioinformatics, 35, 5382–5384.

37. Putman, T.E., Schaper, K., Matentzoglu, N., Rubinetti, V.P., Alquaddoomi, F.S., Cox, C., Caufield, J.H., Elsarboukh, G., Gehrke, S., Hegde, H. et al. (2024) The Monarch Initiative in 2024: an analytic platform integrating phenotypes, genes and diseases across species. Nucleic Acids Res, 52, D938–D949.

38. Chandak, P., Huang, K. and Zitnik, M. (2023) Building a knowledge graph to enable precision medicine. Sci Data, 10, 67.

39. Stear, B.J., Mohseni Ahooyi, T., Simmons, J.A., Kollar, C., Hartman, L., Beigel, K., Lahiri, A., Vasisht, S., Callahan, T.J., Nemarich, C.M. et al. (2024) Petagraph: A large-scale unifying knowledge graph framework for integrating biomolecular and biomedical data. Sci Data, 11, 1338.

40. Himmelstein, D.S., Lizee, A., Hessler, C., Brueggeman, L., Chen, S.L., Hadley, D., Green, A., Khankhanian, P. and Baranzini, S.E. (2017) Systematic integration of biomedical knowledge prioritizes drugs for repurposing. Elife, 6.

41. Schafer, H., Idrissi-Yaghir, A., Arzideh, K., Damm, H., Pakull, T.M.G., Schmidt, C.S., Bahn, M., Lodde, G., Livingstone, E., Schadendorf, D. et al. (2024) BioKGrapher: Initial evaluation of automated knowledge graph construction from biomedical literature. Comput Struct Biotechnol J, 24, 639–660.

42. Zhang, Y., Sui, X., Pan, F., Yu, K.X., Li, K.Q., Tian, S.B., Erdengasileng, A., Han, Q., Wang, W.J., Wang, J.A. et al. (2025) A comprehensive large-scale biomedical knowledge graph for AI-powered data-driven biomedical research. Nat Mach Intell, 7, 602–614.

43. Unni, D.R., Moxon, S.A.T., Bada, M., Brush, M., Bruskiewich, R., Caufield, J.H., Clemons, P.A., Dancik, V., Dumontier, M., Fecho, K. et al. (2022) Biolink Model: A universal schema for knowledge graphs in clinical, biomedical, and translational science. Clin Transl Sci, 15, 1848–1855.

44. Callahan, T.J., Tripodi, I.J., Stefanski, A.L., Cappelletti, L., Taneja, S.B., Wyrwa, J.M., Casiraghi, E., Matentzoglu, N.A., Reese, J., Silverstein, J.C. et al. (2024) An open source knowledge graph ecosystem for the life sciences. Sci Data, 11, 363.

45. Wilkinson, M.D., Dumontier, M., Aalbersberg, I.J., Appleton, G., Axton, M., Baak, A., Blomberg, N., Boiten, J.W., da Silva Santos, L.B., Bourne, P.E., et al. (2016) The FAIR Guiding Principles for scientific data management and stewardship. Sci Data, 3, 160018.

46. Caufield, J.H., Putman, T., Schaper, K., Unni, D.R., Hegde, H., Callahan, T.J., Cappelletti, L., Moxon, S.A.T., Ravanmehr, V., Carbon, S. et al. (2023) KG-Hub-building and exchanging biological knowledge graphs. Bioinformatics, 39.

47. Aevermann, B., Zhang, Y., Novotny, M., Keshk, M., Bakken, T., Miller, J., Hodge, R., Lelieveldt, B., Lein, E. and Scheuermann, R.H. (2021) A machine learning method for the discovery of minimum marker gene combinations for cell type identification from single-cell RNA sequencing. Genome Res, 31, 1767–1780.

48. Liu, A., Peng, B., Pankajam, A.V., Duong, T.E., Pryhuber, G., Scheuermann, R.H. and Zhang, Y. (2024) Discovery of optimal cell type classification marker genes from single cell RNA sequencing data. BMC Methods, 1.

49. Zhang, Y., Aevermann, B.D., Bakken, T.E., Miller, J.A., Hodge, R.D., Lein, E.S. and Scheuermann, R.H. (2021) FR-Match: robust matching of cell type clusters from single cell RNA sequencing data using the Friedman-Rafsky non-parametric test. Brief Bioinform, 22.

50. Buniello, A., Suveges, D., Cruz-Castillo, C., Llinares, M.B., Cornu, H., Lopez, I., Tsukanov, K., Roldan-Romero, J.M., Mehta, C., Fumis, L. et al. (2025) Open Targets Platform: facilitating therapeutic hypotheses building in drug discovery. Nucleic Acids Res, 53, D1467–D1475.

51. Jackson, R., Matentzoglu, N., Overton, J.A., Vita, R., Balhoff, J.P., Buttigieg, P.L., Carbon, S., Courtot, M., Diehl, A.D., Dooley, D.M. et al. (2021) OBO Foundry in 2021: operationalizing open data principles to evaluate ontologies. Database (Oxford), 2021.

52. Moxon, S.A.T., Solbrig, H., Harris, N.L., Kalita, P., Miller, M.A., Patil, S., Schaper, K., Bizon, C., Caufield, J.H., Cuesta, S.C. et al. (2026) LinkML: an open data modeling framework. Gigascience, 15.

53. Osumi-Sutherland, D., Xu, C., Keays, M., Levine, A.P., Kharchenko, P.V., Regev, A., Lein, E. and Teichmann, S.A. (2021) Cell type ontologies of the Human Cell Atlas. Nat Cell Biol, 23, 1129–1135.

54. Vasilevsky, N.A., Toro, S., Matentzoglu, N., Flack, J.E., Mullen, K.R., Hegde, H., Gehrke, S., Whetzel, P.L., Shwetar, Y., Harris, N.L. et al. (2026) Mondo: integrating disease terminology across communities. Genetics, 232.

55. Mungall, C.J., Torniai, C., Gkoutos, G.V., Lewis, S.E. and Haendel, M.A. (2012) Uberon, an integrative multi-species anatomy ontology. Genome Biol, 13, R5.

56. Brown, G.R., Hem, V., Katz, K.S., Ovetsky, M., Wallin, C., Ermolaeva, O., Tolstoy, I., Tatusova, T., Pruitt, K.D., Maglott, D.R. et al. (2015) Gene: a gene-centered information resource at NCBI. Nucleic Acids Res, 43, D36–42.

57. UniProt, C. (2025) UniProt: the Universal Protein Knowledgebase in 2025. Nucleic Acids Res, 53, D609–D617.

58. Zdrazil, B., Felix, E., Hunter, F., Manners, E.J., Blackshaw, J., Corbett, S., de Veij, M., Ioannidis, H., Lopez, D.M., Mosquera, J.F., et al. (2024) The ChEMBL Database in 2023: a drug discovery platform spanning multiple bioactivity data types and time periods. Nucleic Acids Res, 52, D1180–D1192.

59. Smith, B., Ceusters, W., Klagges, B., Kohler, J., Kumar, A., Lomax, J., Mungall, C., Neuhaus, F., Rector, A.L. and Rosse, C. (2005) Relations in biomedical ontologies. Genome Biol, 6, R46.

60. Hu, J., Peng, B., Pankajam, A.V., Xu, B., Deshpande, V.A., Bueckle, A., Herr, B.W., 2nd, Borner, K., Dupont, C., Scheuermann, R.H. et al. (2025) Benchmarking single cell transcriptome matching methods for incremental growth of reference atlases. bioRxiv.

61. Di Tommaso, P., Chatzou, M., Floden, E.W., Barja, P.P., Palumbo, E. and Notredame, C. (2017) Nextflow enables reproducible computational workflows. Nat Biotechnol, 35, 316–319.

62. Jorstad, N.L., Close, J., Johansen, N., Yanny, A.M., Barkan, E.R., Travaglini, K.J., Bertagnolli, D., Campos, J., Casper, T., Crichton, K. et al. (2023) Transcriptomic cytoarchitecture reveals principles of human neocortex organization. Science, 382, eadf6812.

63. Rozenblatt-Rosen, O., Regev, A., Oberdoerffer, P., Nawy, T., Hupalowska, A., Rood, J.E., Ashenberg, O., Cerami, E., Coffey, R.J., Demir, E. et al. (2020) The Human Tumor Atlas Network: Charting Tumor Transitions across Space and Time at Single-Cell Resolution. Cell, 181, 236–249.

64. Baker, T.B., S.; Isaac, A.; Miles, A.; Schreiber, G.; Summers, E. (2013) Key choices in the design of Simple Knowledge Organization System (SKOS). Journal of Web Semantics, 20, 35–49.

65. Mays, A.D. (2026) Elements of Style in Biomedical Workflow Creation in the era of Agentic AI.

66. Scheuermann, R.H.D.-M., A.; Diller, M.; LeClair, R.; Spear, W.; Zhang, Y. (2026) A trustworthy data-driven biomedical knowledge base of cell phenotypes for the National Library of Medicine.

67. Xu, C., Prete, M., Webb, S., Jardine, L., Stewart, B.J., Hoo, R., He, P., Meyer, K.B. and Teichmann, S.A. (2023) Automatic cell-type harmonization and integration across Human Cell Atlas datasets. Cell, 186, 5876–5891 e5820.

68. Plasschaert, L.W., Zilionis, R., Choo-Wing, R., Savova, V., Knehr, J., Roma, G., Klein, A.M. and Jaffe, A.B. (2018) A single-cell atlas of the airway epithelium reveals the CFTR-rich pulmonary ionocyte. Nature, 560, 377–381.

69. Montoro, D.T., Haber, A.L., Biton, M., Vinarsky, V., Lin, B., Birket, S.E., Yuan, F., Chen, S., Leung, H.M., Villoria, J. et al. (2018) A revised airway epithelial hierarchy includes CFTR-expressing ionocytes. Nature, 560, 319–324.

70. Hegyi, P., Wilschanski, M., Muallem, S., Lukacs, G.L., Sahin-Toth, M., Uc, A., Gray, M.A., Rakonczay, Z., Jr. and Maleth, J. (2016) CFTR: A New Horizon in the Pathomechanism and Treatment of Pancreatitis. Rev Physiol Biochem Pharmacol, 170, 37–66.

71. Ma, L., Roman-Campos, D., Austin, E.D., Eyries, M., Sampson, K.S., Soubrier, F., Germain, M., Tregouet, D.A., Borczuk, A., Rosenzweig, E.B. et al. (2013) A novel channelopathy in pulmonary arterial hypertension. N Engl J Med, 369, 351–361.

72. Armulik, A., Genove, G. and Betsholtz, C. (2011) Pericytes: developmental, physiological, and pathological perspectives, problems, and promises. Dev Cell, 21, 193–215.

73. Marucci, G., Buccioni, M., Ben, D.D., Lambertucci, C., Volpini, R. and Amenta, F. (2021) Efficacy of acetylcholinesterase inhibitors in Alzheimer’s disease. Neuropharmacology, 190, 108352.

74. Karran, E. and De Strooper, B. (2022) The amyloid hypothesis in Alzheimer disease: new insights from new therapeutics. Nat Rev Drug Discov, 21, 306–318.

75. Kocis, P., Tolar, M., Yu, J., Sinko, W., Ray, S., Blennow, K., Fillit, H. and Hey, J.A. (2017) Elucidating the Abeta42 Anti-Aggregation Mechanism of Action of Tramiprosate in Alzheimer’s Disease: Integrating Molecular Analytical Methods, Pharmacokinetic and Clinical Data. CNS Drugs, 31, 495–509.

76. de Oliveira, F.F., Bertolucci, P.H., Chen, E.S. and Smith, M.C. (2014) Brain-penetrating angiotensin-converting enzyme inhibitors and cognitive change in patients with dementia due to Alzheimer’s disease. J Alzheimers Dis, 42 Suppl 3, S321–324.

77. Kiser, P.D. (2022) Retinal pigment epithelium 65 kDa protein (RPE65): An update. Prog Retin Eye Res, 88, 101013.

78. Russell, S., Bennett, J., Wellman, J.A., Chung, D.C., Yu, Z.F., Tillman, A., Wittes, J., Pappas, J., Elci, O., McCague, S. et al. (2017) Efficacy and safety of voretigene neparvovec (AAV2-hRPE65v2) in patients with RPE65-mediated inherited retinal dystrophy: a randomised, controlled, open-label, phase 3 trial. Lancet, 390, 849–860.

79. Bombieri, C., Claustres, M., De Boeck, K., Derichs, N., Dodge, J., Girodon, E., Sermet, I., Schwarz, M., Tzetis, M., Wilschanski, M., et al. (2011) Recommendations for the classification of diseases as CFTR-related disorders. J Cyst Fibros, 10 Suppl 2, S86–102.

80. Guo, Q., Li, H., Gaddam, S.S., Justice, N.J., Robertson, C.S. and Zheng, H. (2012) Amyloid precursor protein revisited: neuron-specific expression and highly stable nature of soluble derivatives. J Biol Chem, 287, 2437–2445.

81. Morrow, D.A., Braunwald, E., Bonaca, M.P., Ameriso, S.F., Dalby, A.J., Fish, M.P., Fox, K.A., Lipka, L.J., Liu, X., Nicolau, J.C. et al. (2012) Vorapaxar in the secondary prevention of atherothrombotic events. N Engl J Med, 366, 1404–1413.

82. Islam, M.M.X., B.; Pankajam, A. V.; Zweibaum, D.; Deslattes Mays, A.; Spear, W.; Leclair, R.; Mondal, A. M.; Scheuermann, R. H. (2026) Agentic AI for identification of drug-repurposing topological structures in the NLM-CKN.

83. Rotenberg, N.H., Leaman, R., Islamaj, R., Kuivaniemi, H., Tromp, G., Fluharty, B., Richardson, S., Eastwood, C., Diller, M., Xu, B. et al. (2026) Cell phenotypes in the biomedical literature: a systematic analysis and text mining corpus. bioRxiv.

