## Supplementary Figure for "A trustworthy semantic knowledge network linking data-driven cell phenotypes to diseases and drugs"

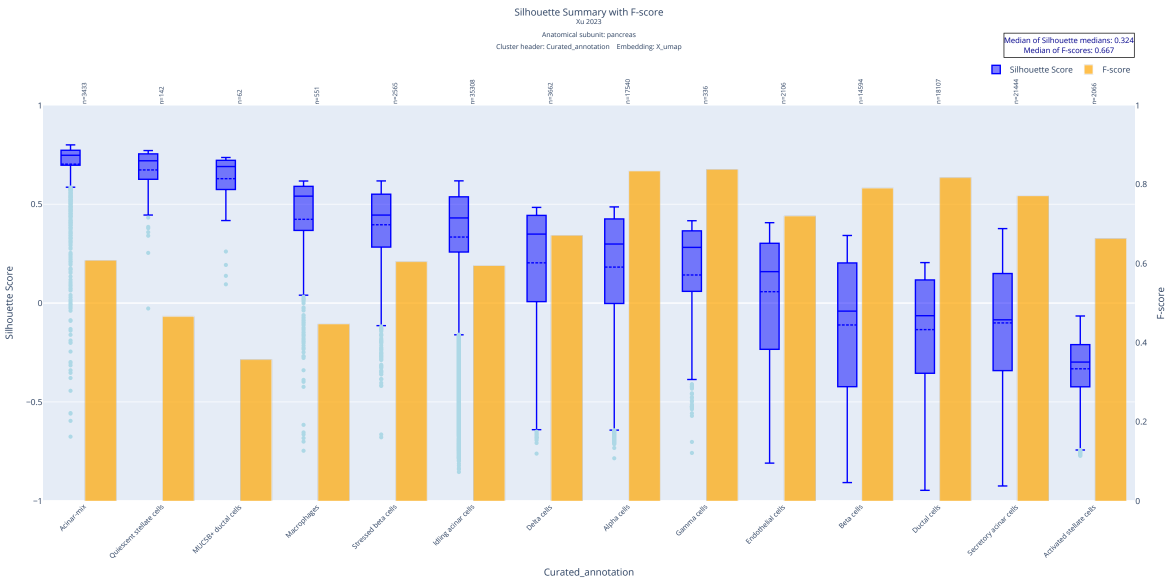


**Supplementary Figure S1.** Quality control metrics for a human pancreas dataset "Xu et al (2023) Cell – 'Pancreas' for the pancreas". The dataset was processed using the `sc-nsforest-qc-nf` workflow with the "curated_annotation" `obs` cell set annotations and UMAP embedding coordinates from the .h5ad file. Box-and-whisker plots (blue) display the distribution of silhouette scores for cells within each cell set. Bar plots (orange) display the NS-Forest F-beta scores for marker gene combinations in each cell set across all 61 annotated cell sets in the dataset. The cell sets are ordered in decreasing order of silhouette scores.

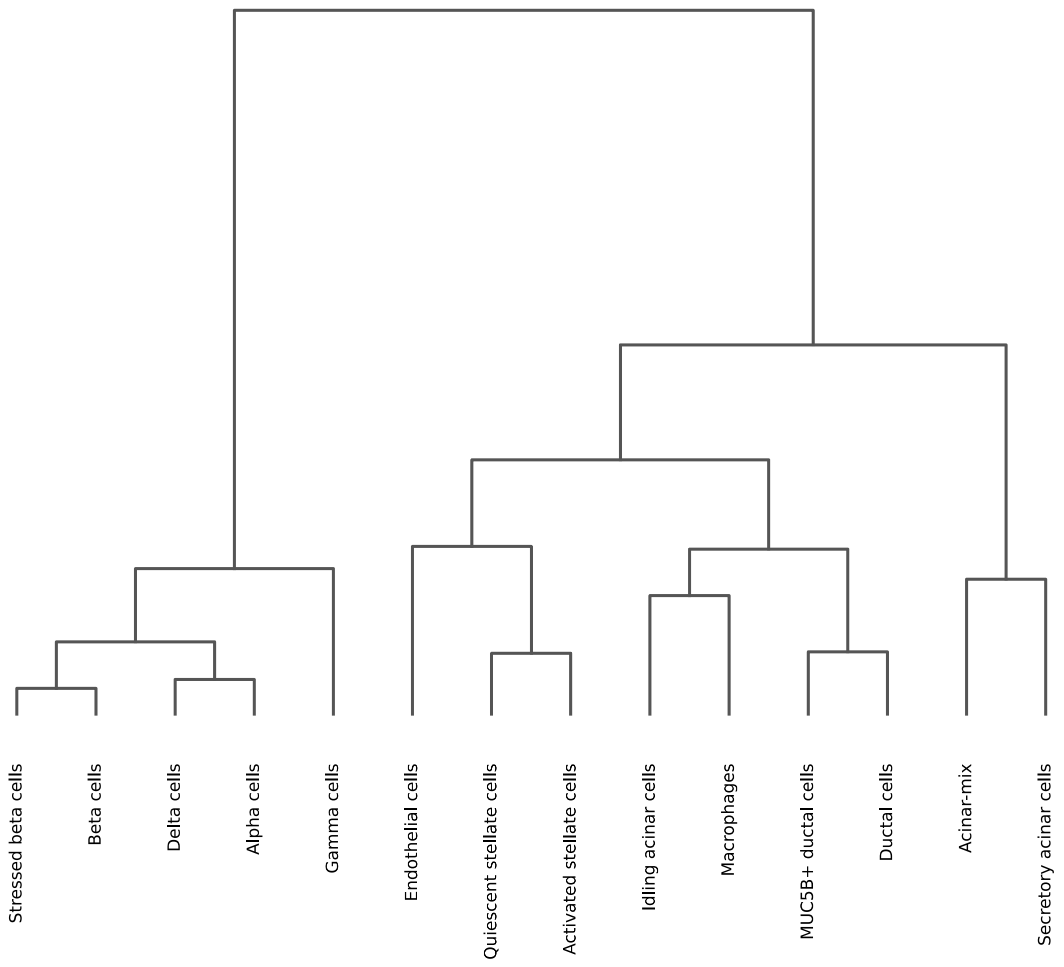


**Supplementary Figure S2.** Hierarchical clustering dendrogram for the "Xu et al (2023) Cell – 'Pancreas' for the pancreas" dataset. The dendrogram is generated during the `sc-nsforest-qc-nf` workflow using the "curated_annotation" cell set annotation in the `.h5ad` `obs` field.

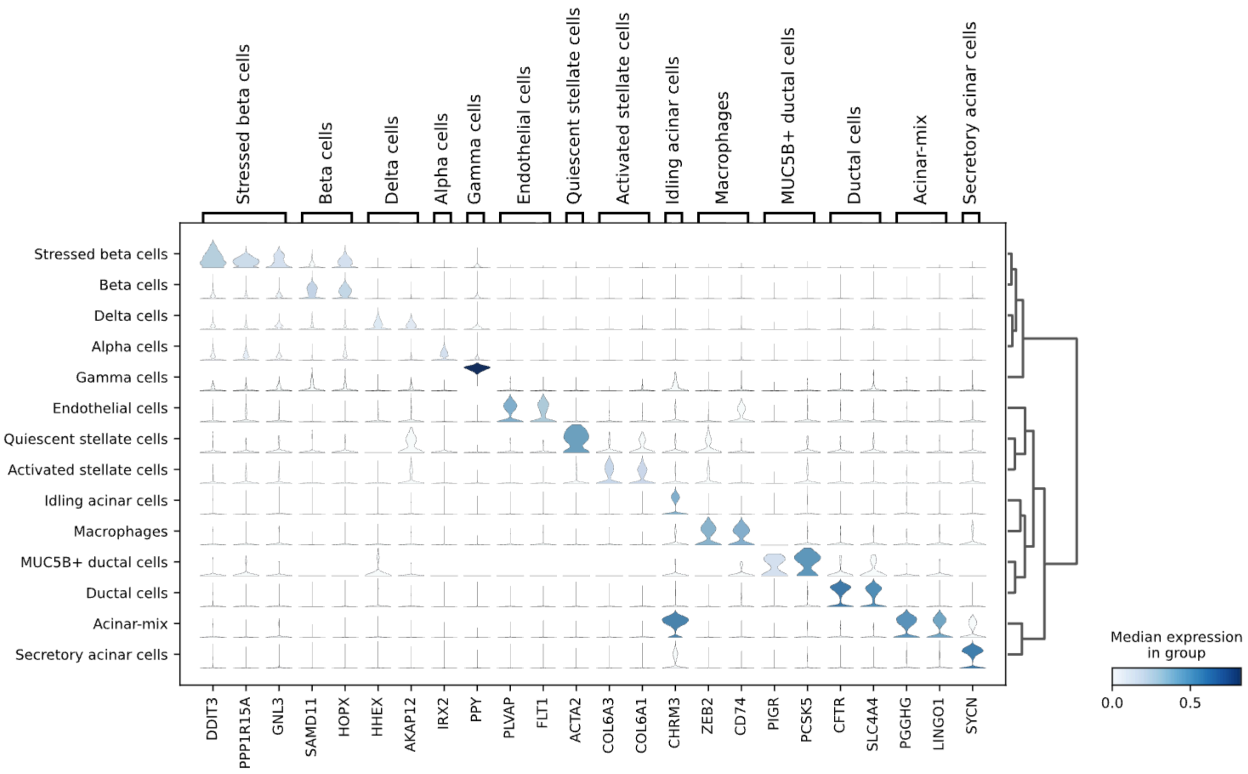


**Supplementary Figure S3.** Marker gene violin expression plot for the "Xu et al (2023) Cell – 'Pancreas' for the pancreas" dataset. The violin plot is generated during the `sc-nsforest-qc-nf` workflow using marker gene and cell set information and expression data from `.h5ad`. The x-axis represents the marker gene combinations for the annotated cell sets. The y-axis represents the annotated cell sets. Ideal markers show a diagonal expression pattern in the plot. The cell set order follows the dendrogram order in Supplementary Figure S2.

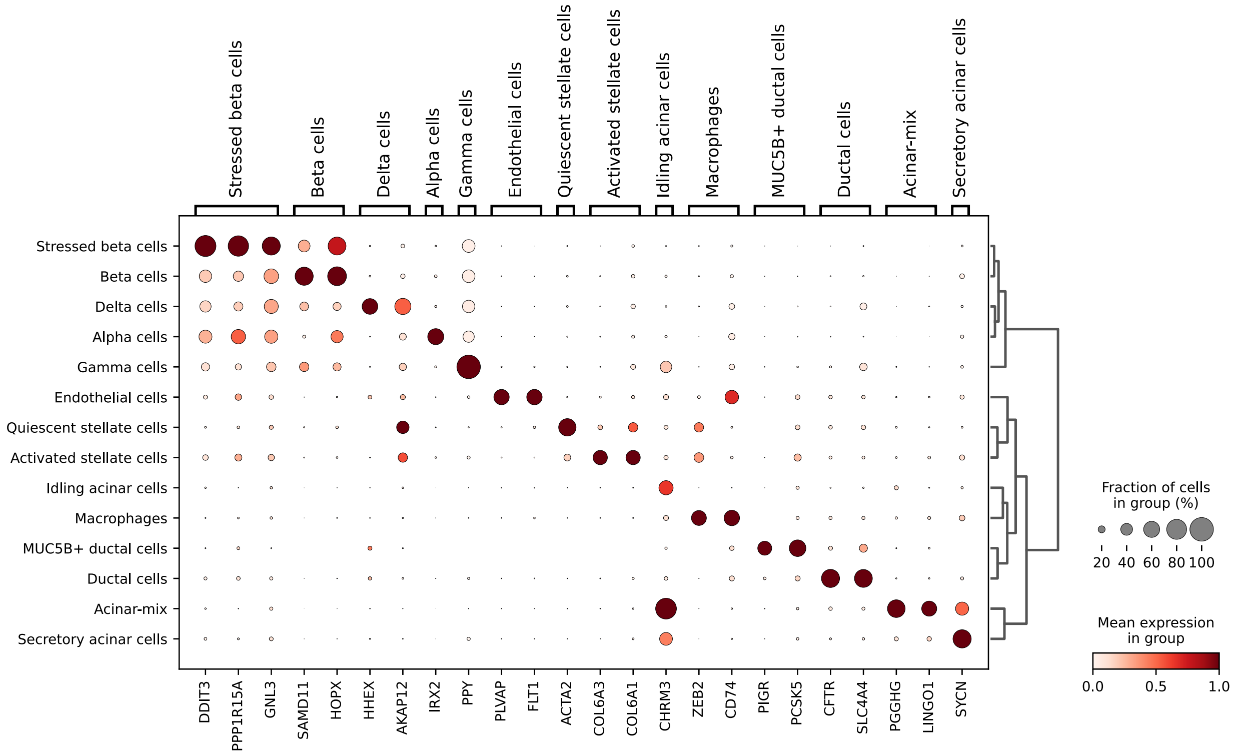


**Supplementary Figure S4.** Marker gene dot expression plot for the "Xu et al (2023) Cell – 'Pancreas' for the pancreas" dataset. The dot plot is generated by the `sc-nsforest-qc-nf` workflow using marker gene and cell set information and expression data from `.h5ad`. The x-axis represents marker gene combinations for the annotated cell sets. The y-axis represents the annotated cell sets. Ideal markers show a diagonal expression pattern in the plot. The cell set order follows the dendrogram order in Supplementary Figure S2.

Class1

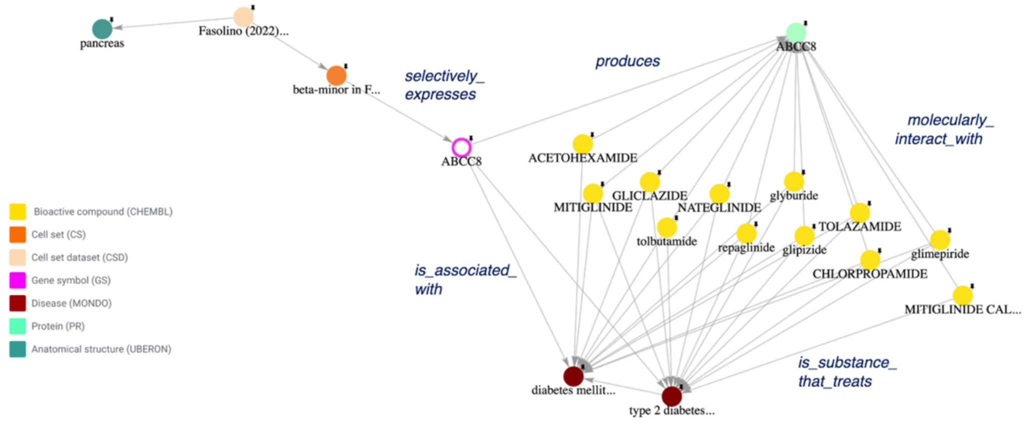


**Supplementary Figure S5.** Example of a Class 1 Big Dipper topology. *ABCC8* links pancreatic β-cell biology and diabetes therapeutics. *ABCC8* (pink node) is selectively expressed in the β-minor cell set (orange node) from the pancreas (green node) Fasolino et al. 2022 cell set dataset (tan node). *ABCC8* is genetically associated with type 2 diabetes and diabetes mellitus diseases (brown nodes). Activating variants of *ABCC8* impair glucose-stimulated insulin secretion, contributing to the development of type 2 diabetes (Tarasov et al., 2008). *ABCC8* encodes the SUR1 protein (cyan node), a subunit of the β-cell ATP-sensitive K+ channel that interacts with multiple bioactive compounds (yellow nodes) used to treat type 2 diabetes and diabetes mellitus. Since *ABCC8* is selectively expressed in pancreatic β-minor cells, the topological structure highlights the potential cellular contribution of β-minor cells to *ABCC8*-associated diabetes pathogenesis.

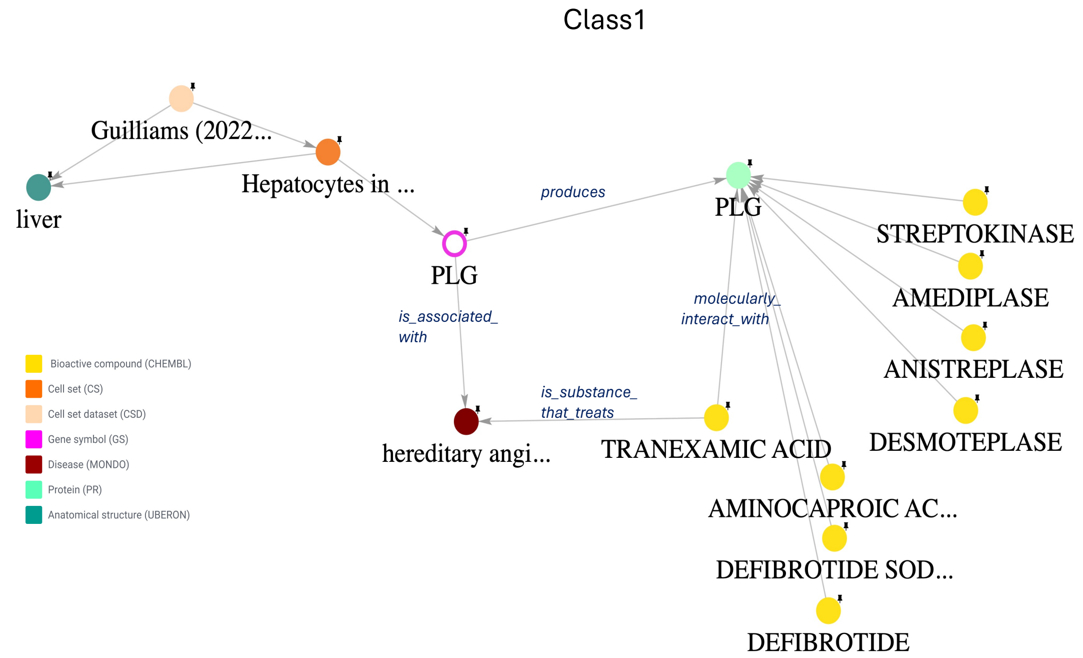


**Supplementary Figure S6.** Example of a Class 1 Big Dipper topology. *PLG* links hepatocyte biology, hereditary angioedema, and therapeutics. *PLG* (pink node) is selectively expressed in the hepatocyte cell set (orange node) from the liver (green node) Guilliams 2022 cell set dataset (tan node). *PLG* is genetically associated with hereditary angioedema (brown node) (Dickeson, Kumar et al. 2022). *PLG* encodes the plasminogen protein (cyan node), the circulating precursor of plasmin, a protease that mediates fibrin degradation and fibrinolysis (Raum, Marcus et al. 1980). PLG molecularly interacts with multiple bioactive compounds (yellow nodes); among them, tranexamic acid is approved for the treatment of hereditary angioedema, while other compounds, including aminocaproic acid and defibrotide/defibrotide sodium, have the potential to treat the disease. In contrast, streptokinase, amediplase, anistreplase, and desmoteplase act in the opposite direction. Plasminogen is produced predominantly by hepatocytes in the liver and secreted into the circulation (Raum, Marcus et al. 1980), consistent with the selective expression of *PLG* in liver hepatocytes observed in NLM-CKN. Together, the hepatocyte-specific expression of *PLG* highlights the potential cellular contribution of hepatocytes to hereditary angioedema.

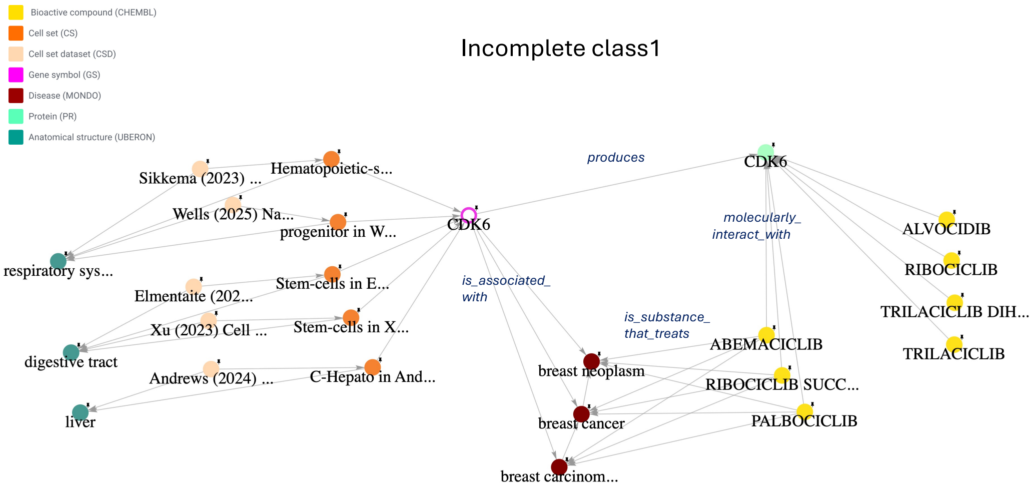


**Supplementary Figure S7.** Example of an incomplete Class 1 Big Dipper topology. *CDK6* links stem cell biology and cancer therapeutics. *CDK6* (pink node) is selectively expressed in multiple progenitor cell sets (orange nodes) in multiple cell set datasets (tan nodes) in multiple tissues (green nodes). *CDK6* is genetically associated with breast cancer and related breast neoplasms (brown nodes). The CDK6 protein (cyan node) is a cyclin-dependent kinase involved in cell cycle progression and cellular proliferation (Nebenfuehr, Kollmann, & Sexl, 2020). CDK4/6 inhibitors (yellow nodes), including abemaciclib, ribociclib, and palbociclib, are used to treat subtypes of breast cancer, whereas other CDK6-interacting compounds shown in the network have different therapeutic indications or are investigational agents. Because breast tissue is not represented in NLM-CKN v1.0, the specific breast cell populations contributing to the *CDK6*-associated breast cancer mechanisms remain to be investigated.

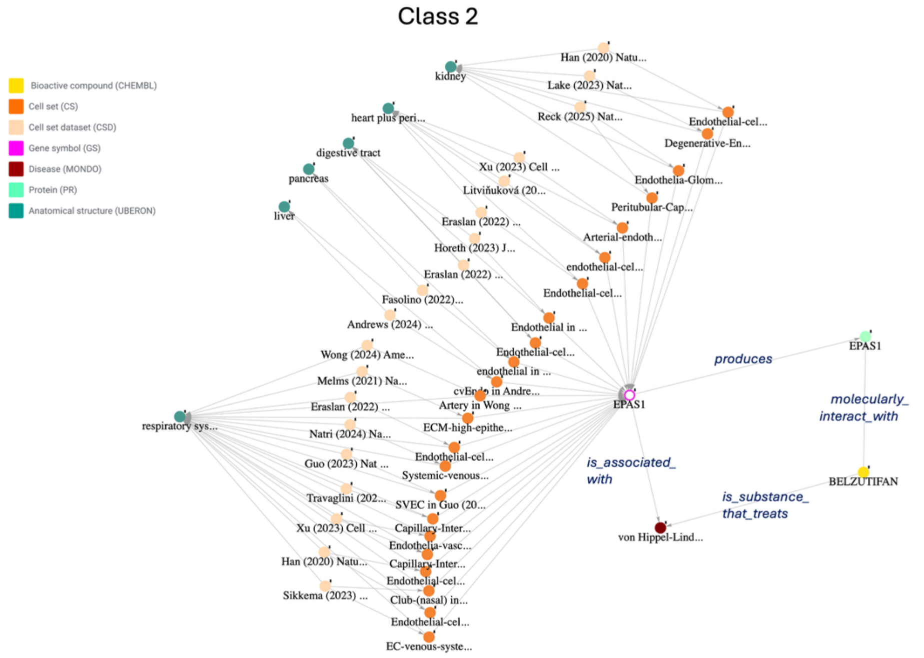


**Supplementary Figure S8.** Example of a Class 2 Big Dipper topology. *EPAS1* links endothelial cell biology, von Hippel–Lindau disease (VHL), and targeted therapy. *EPAS1* (pink node) is selectively expressed in multiple endothelial cell sets (orange nodes) in multiple different cell set datasets (tan nodes) in multiple different tissues (green nodes). VHL disease (brown node) is a rare inherited tumor-predisposition syndrome caused by pathogenic *VHL* variants. Loss of VHL function causes HIF-2α accumulation and activation of hypoxia-responsive genes that promote angiogenesis and tumor growth (Jonasch et al., 2021). *EPAS1* encodes endothelial PAS domain protein 1 (cyan node), also known as hypoxia-inducible factor-2α (HIF-2α), a transcription factor that regulates hypoxia-responsive genes. Although *EPAS1* is not genetically associated with VHL disease, belzutifan (yellow node) directly targets HIF-2α and inhibits HIF-2α-dependent transcription, providing a therapeutic connection between *EPAS1* and VHL disease. Since *EPAS1* is selectively expressed in endothelial cells across multiple tissues in NLM-CKN, this expression pattern highlights the potential cellular context of *EPAS1*-targeted therapy.

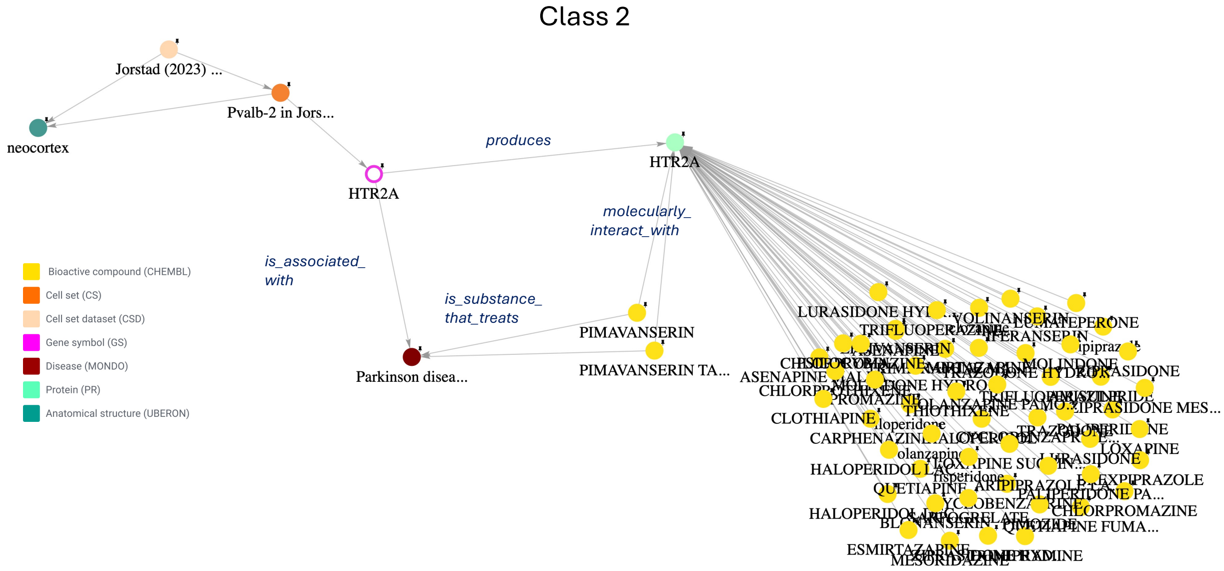


**Supplementary Figure S9**. Example of a Class 2 Big Dipper topology. HTR2A-targeted therapy for Parkinson disease psychosis. *HTR2A* (pink node) is selectively expressed in the Pvalb-2 cell set (orange node) from the Jorstad 2023 cell set dataset (tan node), derived from the brain neocortex (green node). *HTR2A* encodes the serotonin 5-HT2A receptor (cyan node), a GPCR targeted by numerous neuroactive compounds (yellow nodes). Although *HTR2A* is associated with Parkinson disease (brown node), it is not an established causal gene for the disease. Inhibition of 5-HT2A signaling provides a therapeutic mechanism for the treatment of Parkinson disease psychosis; in particular, pimavanserin is a selective 5-HT2A inverse agonist used to treat hallucinations and delusions associated with Parkinson disease psychosis (Hawkins & Berman, 2017). The selective expression of *HTR2A* in the Pvalb-2 cell set from the human neocortex highlights the potential cellular context for *HTR2A*-targeted pharmacological effects.

### References

Hawkins, T., & Berman, B. D. (2017). Pimavanserin: A novel therapeutic option for Parkinson disease psychosis. *Neurol Clin Pract, 7*(2), 157–162. doi:10.1212/CPJ.0000000000000342

Jonasch, E., Donskov, F., Iliopoulos, O., Rathmell, W. K., Narayan, V. K., Maughan, B. L., . . . Investigators, M. K. (2021). Belzutifan for Renal Cell Carcinoma in von Hippel-Lindau Disease. *N Engl J Med, 385*(22), 2036–2046. doi:10.1056/NEJMoa2103425

Nebenfuehr, S., Kollmann, K., & Sexl, V. (2020). The role of CDK6 in cancer. *Int J Cancer, 147*(11), 2988–2995. doi:10.1002/ijc.33054

Tarasov, A. I., Nicolson, T. J., Riveline, J. P., Taneja, T. K., Baldwin, S. A., Baldwin, J. M., . . . Rutter, G. A. (2008). A rare mutation in ABCC8/SUR1 leading to altered ATP-sensitive K+ channel activity and beta-cell glucose sensing is associated with type 2 diabetes in adults. *Diabetes, 57*(6), 1595–1604. doi:10.2337/db07-1547
